# Derivatization of the non-ribosomal peptide pyrrolizixenamide using NRPS engineering

**DOI:** 10.64898/2026.07.12.738029

**Authors:** Juliana Effert, Andrea Calderari, Susanna Kremer, Kira J. Weissman, Helge B. Bode

**Affiliations:** Max-Planck-Institute for Terrestrial Microbiology, Department of Natural Products in Organismic Interactions, 35043, Marburg, Germany; Université de Lorraine, CNRS, IMoPA, F-54000 Nancy, France; Center for Synthetic Microbiology (SYNMIKRO), Phillips University Marburg, 35043 Marburg, Germany; Department of Chemistry, Phillips University Marburg, 35043 Marburg, Germany

**Keywords:** natural products, pyrrolizidine alkaloids, nonribosomal peptides, NRPS engineering

## Abstract

Pyrrolizidine alkaloids (PA) are well-known and widespread natural products from plants, which have also been identified in several different bacteria. In the latter case, the core structure is constructed by a non-ribosomal peptide synthetase (NRPS), which then undergoes oxidative ring contraction catalyzed by a Baeyer-Villiger monooxygenase. By deploying various NRPS engineering strategies, we have successfully generated five novel peptides carrying the unusual PA moiety at their C-terminus. Nonetheless, efforts to obtain a larger library of PAs were unsuccessful. Combined computational modelling and docking experiments suggest that this failure stems from the strict specificity of the thioesterase (TE) domain at the end of the NRPS, which discriminates against peptides carrying more than two amino acids. Our work thus suggests protein design strategies by which this intrinsic limitation to NRPS engineering may be overcome in future.

## Introduction

Natural products (NP) serve as invaluable starting points for the development of active compounds with novel or enhanced bioactivity for therapeutic applications. Among the diverse classes of NPs, nonribosomal peptides (NRPs) stand out due to their broad distribution and abundance in bacteria and fungi,^[1]^ as well as their remarkable structural diversity.^[2,3]^ NRPs vary significantly in size, ranging from 1–25 amino acids, and exhibit diverse forms, including linear, cyclic and branched structures. Collectively, they incorporate more than 300 non-proteinogenic amino acids, which endows them with a wide range of distinct physicochemical properties relative to canonical peptides.^[3–5]^

A common feature of all NRPs is their biosynthesis by nonribosomal peptide synthetases (NRPS). These megaenzymes are organized in modular fashion, with each module responsible for incorporating a specific building block into the growing peptide.^[3]^ Each NRPS module activates an amino acid and couples the monomer to the elongating chain. For this, minimal NRPS modules comprise three essential domains: adenylation (A), thiolation (T), and condensation (C). The A domain is responsible for selecting and activating the amino acid building block, and then loading it onto the 4′-phosphopantetheine (Ppant) prosthetic group of the T domain. The T domain, which comprises a structurally conserved four α-helix bundle,^[6]^ delivers the amino acid to the C domain, which catalyzes the condensation reaction between the new building block and the peptide intermediate. Thus, the elongating peptide grows from its *C*-to its *N*-terminus as it is shuttled from one module to the next. Optional domains that structurally diversify the chains include epimerization (E), heterocyclization (HC), *N*-methylation (MT) and oxidation (Ox) domains. When the peptide chain reaches the last module, an offloading domain (in most cases a thioesterase (TE)) releases the oligopeptide from the NRPS, either as a linear or cyclic product.^[3]^

The intrinsic modularity of NRPS has inspired the development of engineering strategies to create hybrids of selected NRPS domains and modules,^[7]^ which when functional, give rise to modified NRPs whose bioactivities can be evaluated. Among the most successful approaches is the eXchange Unit (XU) concept,^[8]^ which relies on A-T-C or A-T-C/E domain sets as the basic engineering units. On a practical level, XUs must be fused within the C-A linker at a well conserved WNATE sequence motif. This strategy has notably enabled the assembly of up to five XUs from different NRPS systems, and yielded functional *de novo* NRPS that synthesized the desired peptides in reasonable yields.^[8]^

In more recent work, we developed the concept of eXchange Units between T domains (XUT),^[9]^ which relies on an evolutionarily inspired fusion point within the T domains – an FFxxGGxS motif situated at the *C*-terminal end of loop 1. Successful NRPS fusions have been engineered both upstream of the two phenylalanine (XUT^III^) and the two glycine residues (XUT^IV^), respectively. An additional fusion site flagged by a conserved tyrosine has been identified in the A-T linker (XUT^I^). Use of the resulting T-C-A (XUT^I^) and T_1/2_-C-A-T_1/2_ units (XUT^III^ and XUT^IV^)^[9]^ for NRPS engineering has already facilitated the assembly of NRPS fragments spanning a wide range of protein similarity levels and GC contents of the underlying NRPS-encoding genes,^[9,10]^ as well as the engineering of functional NRPS-polyketide synthase (PKS) hybrids.^[11]^

In this work, we sought to apply these strategies to the structural diversification of pyrrolizixenamide (**1a**, Fig. 1)^[12]^ a bacterial pyrrolizidine alkaloid (PA).^[13]^ PAs are widespread natural products which are produced by plants and microorganisms. They are defined by their pyrrolizidine core structure, which is a bicyclic aliphatic hydrocarbon consisting of two ortho-fused five-membered rings with a bridgehead nitrogen. Investigation of over 40 identified microbial PAs has revealed that the backbone is formed by a common route involving an NRPS and a Baeyer-Villiger monooxygenase (BVMO).^[12,14]^

**Figure 1.**
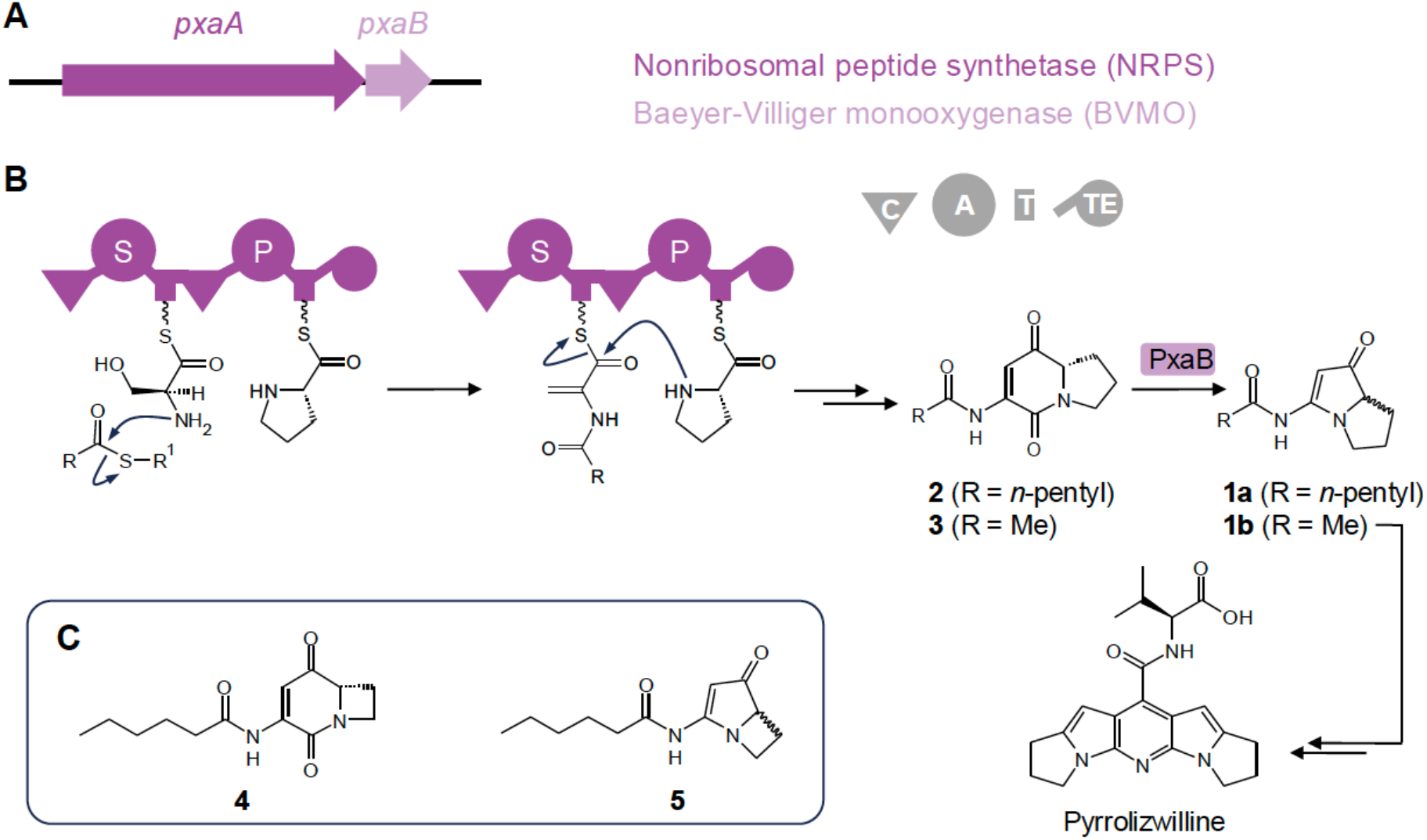
Biosynthesis of pyrrolizixenamides in *Xenorhabdus*. **A**) The BGC *pxaAB* from *X. stockiae* encodes the NRPS PxaA and the BVMO PxaB. **B**) Schematic overview of the biosynthesis of pyrrolizixenamide (**1a**) in *X. stockiae*^[12]^ and pyrrolizwilline^[16]^ in *X. hominickii* via **1b** as key intermediate. The NRPS PxaA and XhpA gives rise to the indolizidine derivatives **2** and **3**, respectively, which are then transformed by the BVMO PxaB, leading to **1a** and **1b**; R^1^ = CoA. **C**) Co-expression of PxaA and AzeJ leads to the production of **4**, which in turn is converted to **5** by PxaB.^[17]^ Key to NRPS domains: C, condensation; A, adenylation; T, thiolation; TE, thioesterase. The amino acid specificities of the A domains in B are shown using the one-letter code.

In the entomopathogenic bacterium *Xenorhabdus stockiae*, the biosynthetic pathway leading to the production of **1a** is encoded by the *pxaAB* biosynthetic gene cluster (BGC) (Fig. 1).^[12]^ Specifically, the bimodular NRPS PxaA (domain composition: C_starter_-A^Ser^-T-C_mod_-A^Pro^-T-TE), extends *N*-acylated serine (catalyzed by the C_starter_ domain) with proline after dehydration of serine by the C_mod_ domain (C_mod_ domains show sequence similarity to ^D^C_L_ domains that condense D and L-amino acids, but no upstream E domain is present to generate a D-amino acid), followed by TE-mediated cyclization to give a 5,6-bicyclic indolizidine (**2**) (Fig. 1b). The BVMO PxaB then catalyzes oxygen insertion, leading to a 5,7-bicyclic intermediate. Spontaneous ring contraction and decarboxylation yield the final product pyrrolizixenamide (**1a**)^[12]^ but may also result in carbamate formation, as proposed for brabantamide biosynthesis catalyzed by the PxaB homolog BraC.^[15]^ The biosynthesis of the pyrrolizixenamide-derived pseudodimer pyrrolizwilline follows an analogous mechanism, wherein the NRPS XhpA assembles an analogue of **2** bearing an acetyl side chain (**3**), which is then further processed by the BVMO XhpB to yield **1b** as key intermediate.^[16]^

Previously, we successfully generated PA variants in *E. coli* by co-expressing the azetidine-2-carboxylic acid (AZE) synthase AzeJ from *P. aeruginosa* alongside Pxa components.^[17]^ In particular, PxaA and AzeJ together yielded **4**, a derivative of **2** incorporating AZE instead of proline. Co-expression of AzeJ with both PxaA and PxaB resulted in production of the first engineered analogue of the azabicyclenes (**5**) (Fig. 1).

Here, we successfully leveraged both XU and XUT NRPS engineering approaches to produce six new PA derivatives in which the acyl residue or the first amino acid were modified. In addition, computational modelling of the TE domain of PxaA identified it as a critical bottleneck to obtaining more complex peptide variants.

## Results and Discussion

Initially, we generated hybrids of the closely related C_6_-acyl specific PxaA from *Xenorhabdus stockiae*^[12]^ and C_2_-acyl specific XhpA from *X. hominickii*^[16]^ based on XU and different XUT variants. This series of experiments revealed that XUT^I^ and XUT^III^ were the most productive fusion sites,^[9]^ while XU^[8]^ worked only in one of the four positions tested (Fig. 2, Fig. S1). As expected, these experiments also confirmed the specificity of the starter C domains for their respective acyl chains. We further showed that PxaA could be split into two fragments using synthetic zippers (SYNZIPs) SZ17/18^[18]^ (Fig. S2a).^[19,20]^ Although the amount of the produced peptide was only 21–89% compared to intact PxaA and the yields strongly depended on the position of the split, this approach might still be useful for future engineering campaigns.

**Figure 2.**
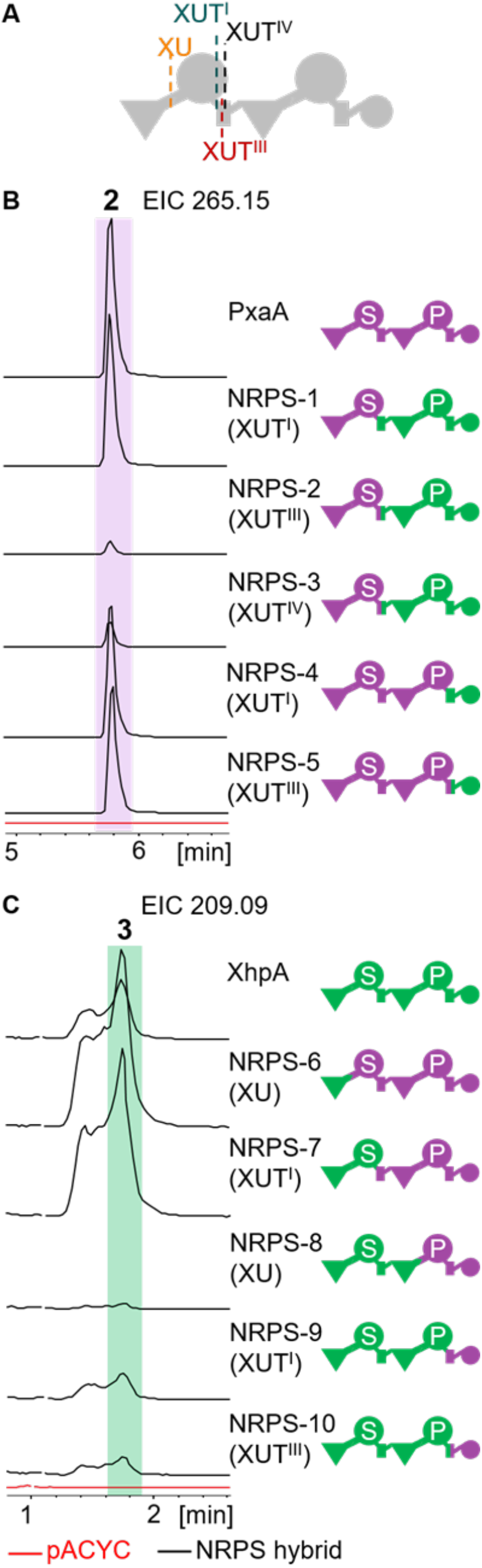
NRPS engineering of C_6_-specific PxaA from *X. stockiae*^[12]^ by exploiting the C_2_-specific XhpA from *X. hominickii*.^[16]^ **A**) Schematic overview of the fusion sites used for NRPS engineering: eXchange Units^[8]^ (XU) and eXchange Units between T domains^[9]^ (XUT^I^, XUT^III^, XUT^IV^). **B**) Engineering of PxaA−XhpA hybrids via the indicated XUT approaches. The EIC of **2** (*m/z* = 265.15 [M+H]^+^) for the respective NRPS hybrid (black) was compared to the empty vector control (red). Peak intensity of the parental PxaA variant was 4 x 10^8^. **C**) Engineering of XhpA−PxaA hybrids via the indicated XUT approaches. The EIC of **3** (*m/z* = 209.09 [M+H]^+^) for the respective NRPS hybrid (black) was compared to the empty vector control (red). Peak intensity of the parental XhpA variant was 1 x 10^7^.The unusual peak form of **3** was previously also observed for its final end product **1b**.^[16]^ All chromatograms are shown relative to production by the original variant shown at the top in **B** and **C**, respectively.

We next tested whether C_starter_ or C_starter_-A domains of the unrelated NRPSs XabA^[21]^ and TxlA^[22]^ producing the depsipeptides xenoamicin and taxlllaid, respectively, were also functional. The expected products were successfully obtained, but at 1–10% of the titers of compound **2** (see Fig. 2) as determined by their peak intensity (Fig. 3, Fig. S3). Additionally, these experiments revealed that Thr is accepted in place of Ser as the first amino acid, with the product serving as a substrate for the PxaB-catalyzed ring contraction (Fig. 3c, Fig. S3). Interestingly, related Thr-containing PAs have already been described from various *Streptomyces* strains,^[14]^ but not yet from proteobacteria.

**Figure 3.**
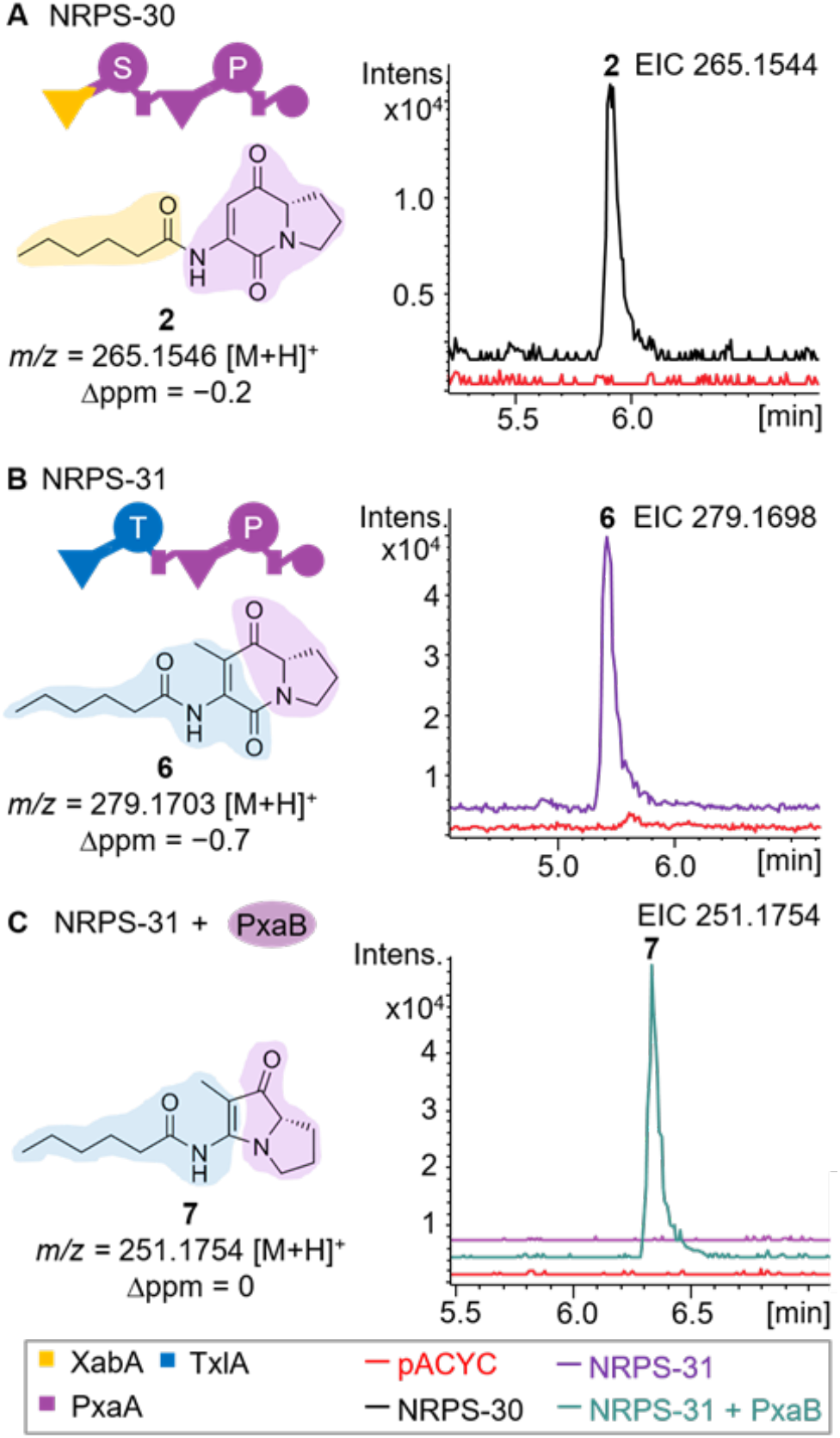
Engineering of functional NRPS hybrids XabA-PxaA and TxlA-PxaA. For detailed HPLC/MS data see Fig. S3. **A**) The C_starter_ domain of the XabA^[21]^ NRPS from *Xenorhabdus* KJ12.1 was introduced into PxaA using the XU concept. The resulting NRPS-30 produced **2** (black chromatogram), as judged by comparison to the empty vector control (red chromatogram). EIC for **2** (*t*_*R*_ = 5.9 min; *m/z* calc’d for C_14_H_20_N_2_O_3_ [M+H]^+^ = 265.1547; obs’d. *m/z* = 265.1544; Δppm = −0.2). **B**) The C_starter_–A didomain of the TxlA^[38]^ NRPS from *X. bovienii* was grafted onto PxaA using the XUT concept. The resulting NRPS-31 was shown to produce **6** (*t*_*R*_ = 5.2 min), as judged by comparison to the empty vector control. EIC for **6** (*t*_*R*_ = 5.2 min; *m/z* calc’d for C_14_H_20_N_2_O_3_ [M+H]^+^ = 265.1547; obs’d. *m/z* = 265.1544; Δppm = −1.1). **C**) The pyrrolizixenamide analogue **7** was produced upon co-expression of NRPS-31 and PxaB and its identity was confirmed by LC-HRMS/MS analysis (*t*_*R*_ = 6.3 min, *m*/*z* calc’d for C_14_H_22_N_2_O_2_ [M+H]^+^ = 251.1754; obs’d. *m/z* = 251.1754; Δppm = 0).

Encouraged by these results, we varied both the acyl-incorporating C_starter_ domain and the first A domain, resulting in hybrids NRPS-63 and -64. This approach yielded the expected C_14_-acyl variant **8** containing Thr, which was accepted for ring contraction when coexpressed with PxaB leading to **9** (Fig. 4b, Fig. S4). Unfortunately, all other generated variants of PxaA (NRPS-34 to -48) and further variants of the initially functional NRPS-30 and -31 (see Fig. 3) (NRPS-49 to -62), led neither to the expected products nor to any other peptide variants (Fig. 5, Fig. S5).

**Figure 4.**
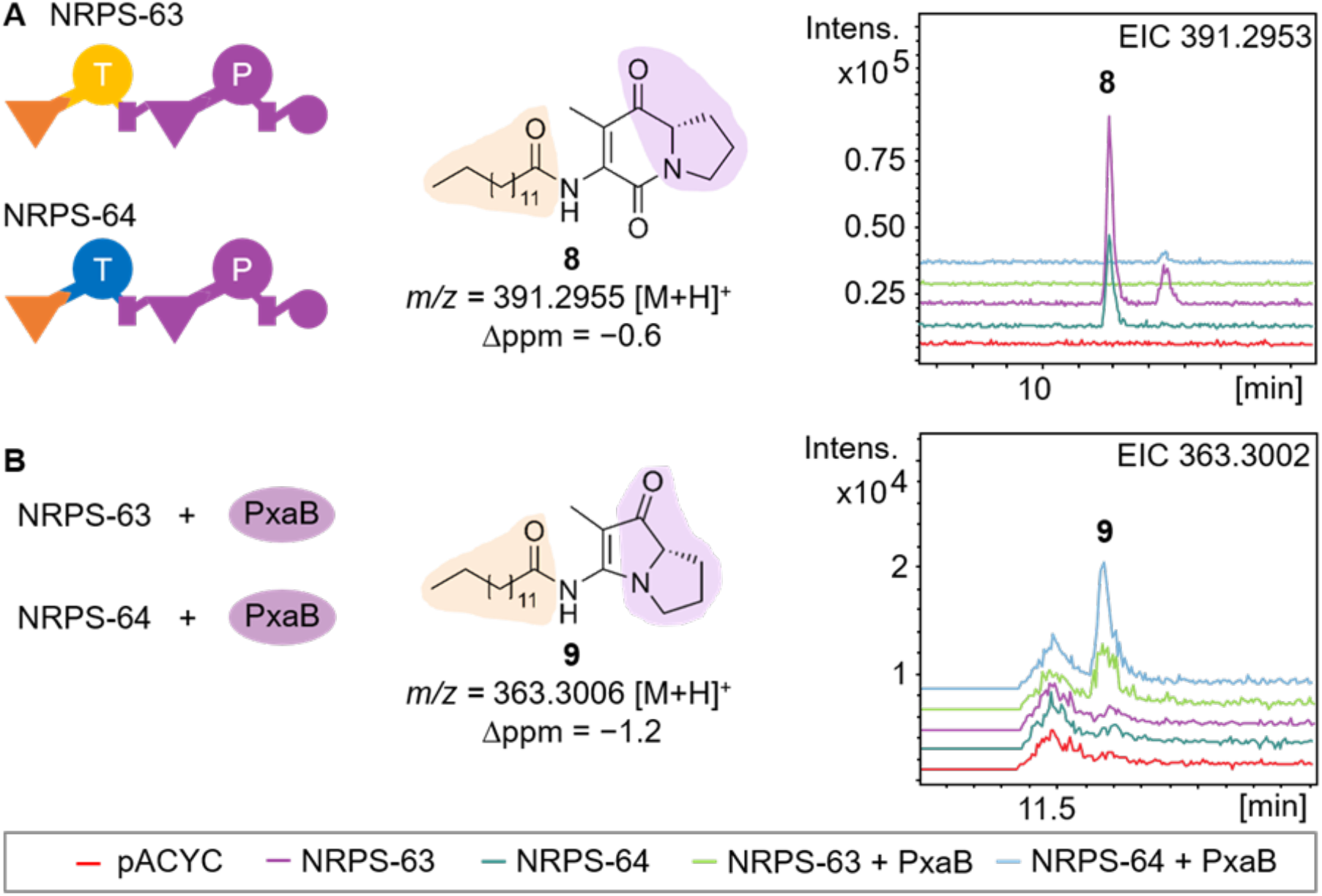
The C_starter_ domain of the XldS^[39]^ from *X. indica* NRPS was introduced into NRPS-30 and NRPS-31 (Fig. S3) using the XU concept, yielding NRPS-63 and NRPS-64, respectively. For detailed HPLC/MS data see Fig. S4. **A**) Both NRPS-63 and NRPS-64 produced **8** (*t*_*R*_ = 10.3 min, *m*/*z* calc’d for C_23_H_38_N_2_O_3_ [M+H]^+^ = 391.2955; obs’d. *m/z* = 391.2953; Δppm = −0.6), based on comparison to the empty vector control (pACYC). **B**) Co-expression of NRPS-63 and NRPS-64 with PxaB, respectively, led to the production of **9** (*t*_*R*_ = 11.6 min, *m*/*z* calc’d for C_22_H_38_N_2_O_2_ [M+H]^+^ = 363.3006; obs’d. *m/z* = 363.3002; Δppm = −1.2).

**Figure 5.**
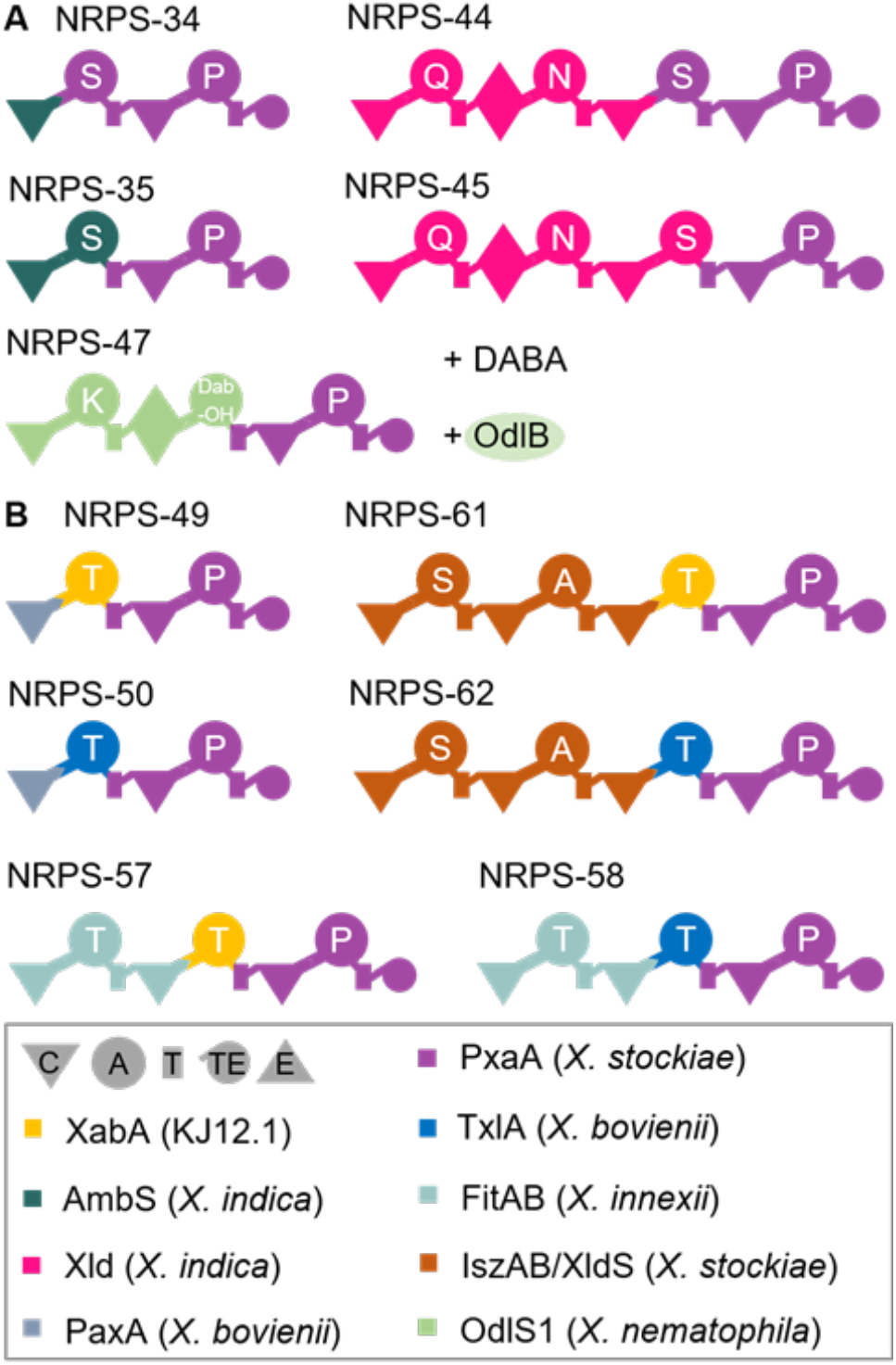
Further engineering of PxaA (**A**) and NRPS-30/NRPS-31 (**B**) resulted in a set of non-functional NRPS hybrids. In the case of NRPS-47, the hydroxylase OdlB^[40]^ from *X. nematophila* was co-expressed in the presence of fed 2,4-diaminobutyric acid (DABA), as the second module in OdlS1 is known to incorporate Dab-OH (3-hydroxy-2,4-diaminobutyric acid). For a complete list of tested NRPS variants see Fig. S5.

To explain the failure to obtain PXA derivatives containing more than two amino acids, we hypothesized that the TE domain was highly specific for acylated dipeptides. To address this issue directly, we initially replaced the TE domain with a homologue from the GameXPeptide producing NRPS GxpS, as the GxpS TE exhibits broad substrate flexibility towards both linear and cyclic peptides.^[23,24]^ However, no indolizidine derivative was obtained, but instead linear lipo-dipeptide **10**. This observation provided further evidence that dehydration of the Ser residue does not depend on the TE^[14]^ but the C_mod_ domain,^[25]^ and that the TE is responsible for the cyclization to yield the indolizidine **2** (Fig. 6). Consistent with this idea, both mutational inactivation of the TE domain and its deletion abolished production of the indolizidine **2** and the linear peptide product **10**. Replacement of the original TE domain with a thioester reductase (R) domain from the tilivalline producing NRPS XtvB,^[26,27]^ resulted in the production of **11**, a reduced variant of **2**, suggesting that cyclization could also occur on an aldehyde containing intermediate (Fig. 7).

**Figure 6.**
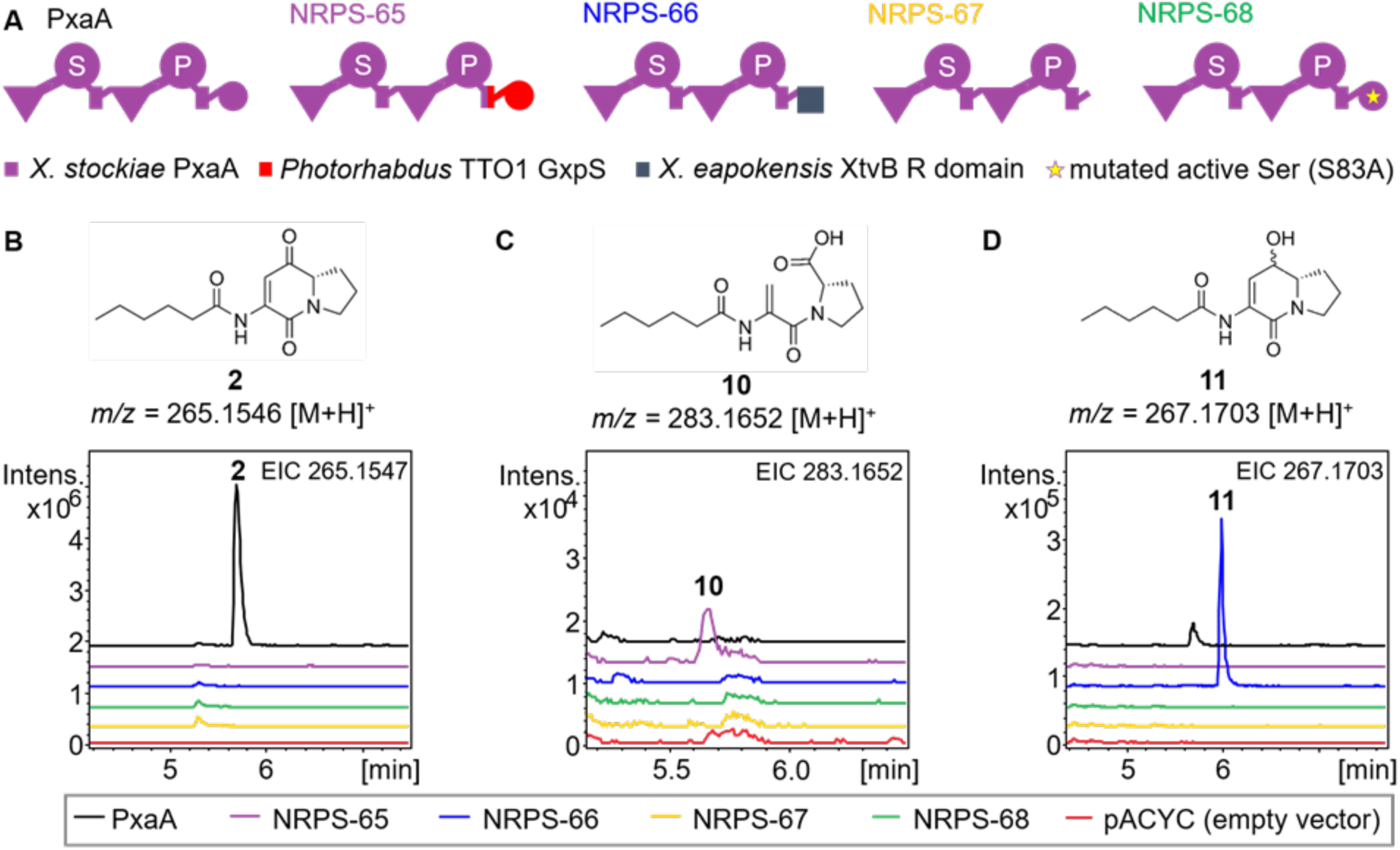
Investigation of the role of the TE domain in PxaA (for MS data acquired on **2, 10** and **11** see Fig. S6). **A**) Schematic overview of the engineered PxaA mutants. In NRPS-65, the TE domain of PxaA was exchanged with the GxpS TE domain from *Photorhabdus* TTO1. In NRPS-66, the PxaA TE domain was exchanged with the R domain of the XtvB NRPS from *X. eapokensis*^[26,27]^. In NRPS-67, the PxaA TE domain was deleted, while in NRPS-68, the active Ser83 of the TE domain was mutated to Ala. **B**) PxaA produces **2** as judged by LC-HRMS/MS analysis. The structure of **2** (*t*_*R*_ = 6.0 min, *m*/*z* calc’d for C_14_H_20_N_2_O_3_ [M+H]^+^ = 265.1547; obs’d *m/z* = 265.1546; Δppm = −0.3) is shown. **C**) NRPS-65 produces **10** as judged by LC-HRMS/MS analysis. The structure of **10** (*t*_*R*_ = 6.2 min, *m*/*z* calc’d for C_14_H_22_N_2_O_4_ [M+H]^+^ = 283.1652; obs’d *m/z* = 283.1652; Δppm = 0) is shown. **D**) NRPS-66 produces **11** as judged by LC-HRMS/MS analysis. The structure of **11** (*t*_*R*_ = 5.6 min, *m*/*z* calc’d for C_14_H_22_N_2_O_3_ [M+H]^+^ = 267.1703; obs’d *m/z* = 267.1703; Δppm = 0) is shown.

**Figure 7.**
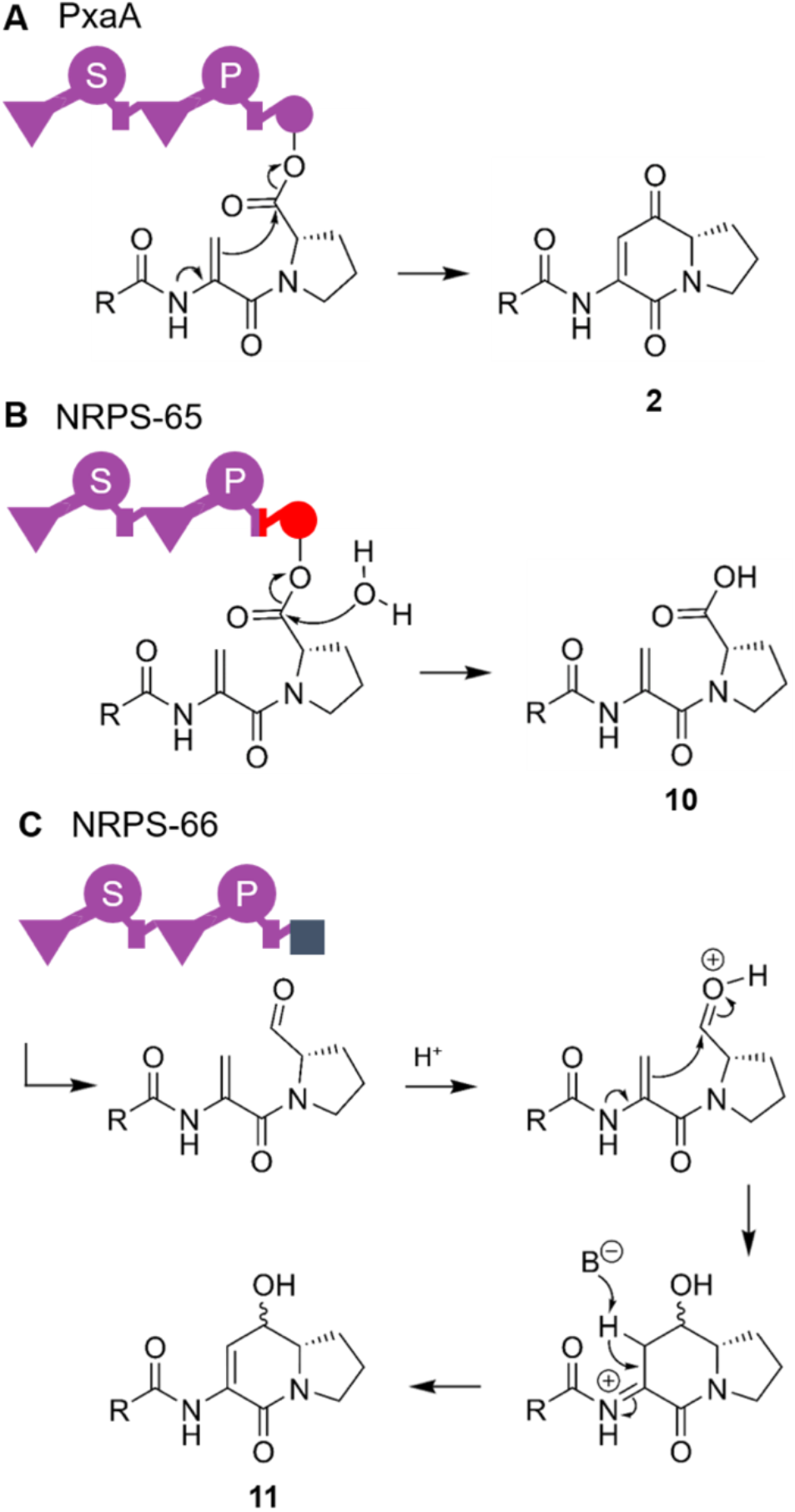
Investigating the offloading of **2** revealed that PxaA TE plays an important role in cyclization. **A**) The PxaA TE catalyzes cyclization to **2** with simultaneous offloading, starting from TE-bound peptide. **B**) The GxpS TE from *Photorhabdus* TTO1 catalyzes coupled hydrolysis and off-loading of TE-tethered **11. C**) The XtvB-R domain from *X. eapokensis* catalyzes offloading of the growing peptide chain while it is attached to the directly upstream T domain. The reactivity of the aldehyde group results in spontaneous cyclization to afford **10**.

To further understand the results obtained from the engineering, we interrogated the PxaA TE acylation half-reaction *in silico* using a combination of AlphaFold and the CB-Dock2 program. For this, we calculated five models of monomeric TE using both AlphaFold2^[28]^ and AlphaFold3^[29]^. We selected one model from each program based on the confidence scores for the active site residues (Fig. S7). Comparison of the two models yielded an RMSD of 0.248 Å, attesting to their close similarity, with the active site triads (Ser83, His256, Asp110; numbering according to the isolated TE domain) essentially superimposable. Notably, the Ser83 rotamers are those seen in the closest structural homologues of the TE.^[30]^ The active site volumes in the selected models were calculated by CB-Dock2^[31]^ to be 405 Å^3^ and 415 Å^3^, respectively.

We then used these two models for docking mimics of the natural substrates **2i** and **5i** into the catalytic cavity. The peptides were derivatized as their *N*-acetylcysteamine thioesters to resemble the natively Ppant-tethered intermediates. We also carried out docking studies with the non-natural substrates **6i** and **8i** of NRPS-31 (Fig. 3) and -63/-64 (Fig. 4), respectively and the proposed intermediates generated by NRPS-44/45 (**12i**), NRPS-57/58 (**13i**) and NRPS-61/62 (**14i**) (see Fig. S5 for NRPS and Fig. S8 for chemical structures used for docking studies), where no product was observed. Despite the strong resemblance of the two AlphaFold models, the results of docking of substrates **2i, 5i** and **6i** into the catalytic cavity were distinct, principally due to differences in the orientation of certain amino acids (Glu134, Trp137, Leu138, Trp193 and Met224) that modify the overall shape of the active site cavity. Globally, we favor the AlphaFold2 model, because both the distance and angle of attack of Ser83 on the thioester better match those reported previously for proteases.^[32]^

Use of this model revealed that substrate mimics **2i** and **6i**, which both incorporate a hexanoyl side chain (volume of substrates calculated with Sanjeevini:^[33]^ 303 Å^3^ and 311 Å^3^, respectively), are oriented similarly within the active site, with the thioester bonds positioned at 2.9 Å and 3.0 Å, respectively, from the catalytic Ser83 (Fig. 8). Ligand **5i** that contains a much shorter acetyl (252 Å^3^), adopts a different but nonetheless catalytically competent position (distance to the catalytic Ser83: 2.9 Å) (Fig. 8). Few specific contacts (i.e. hydrogen bonds) are observed between the TE and the substrates, with recognition rather mediated by hydrophobic and van der Waals interactions and overall shape complementarity (Fig. 8). This binding mode echoes that seen with TEs derived from polyketide synthase (PKS) systems for which there is also a substantial shape-dependence.^[34]^

**Figure 8.**
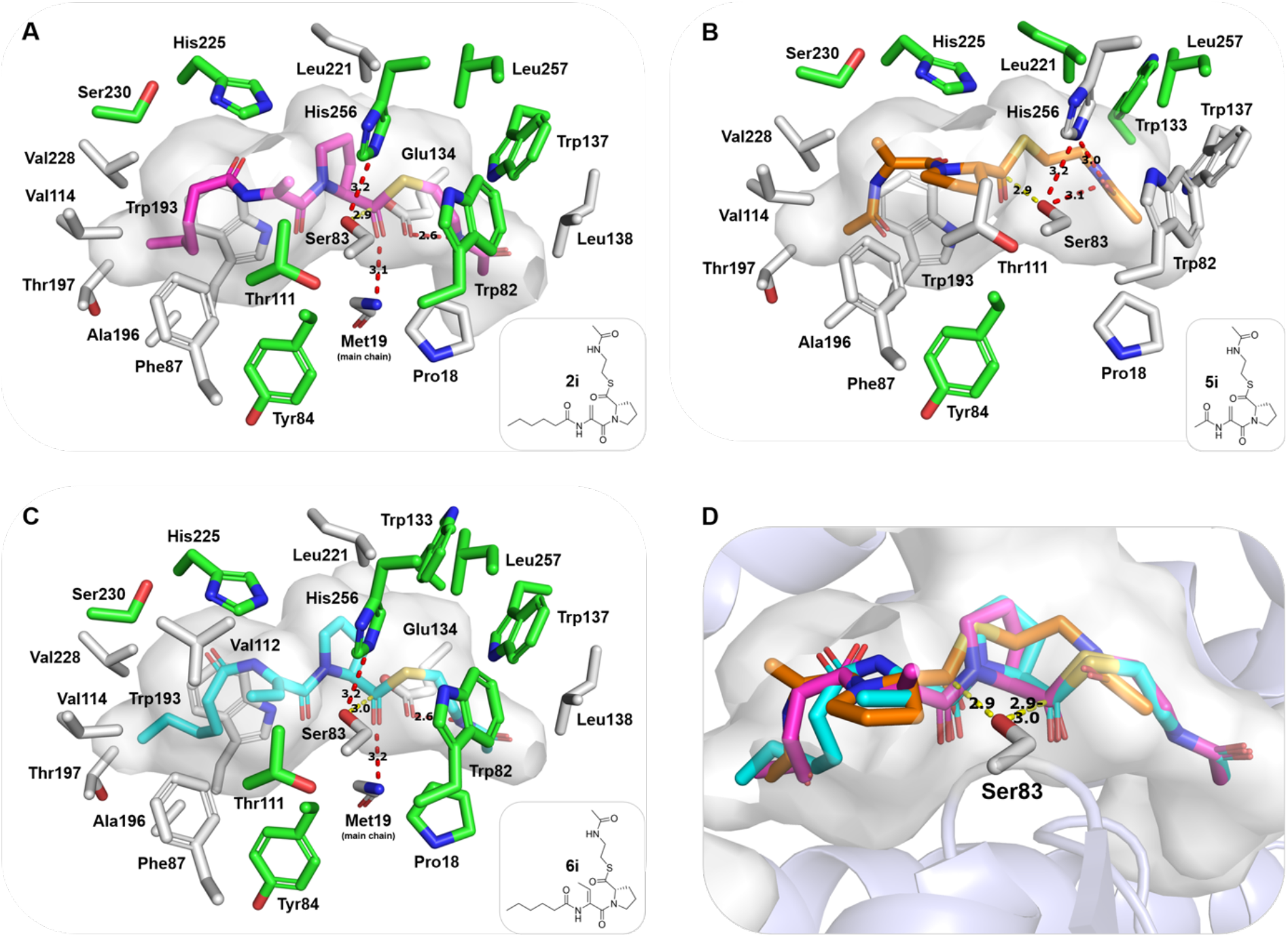
**A–C**) Models for the binding of SNAC-peptide intermediates **2i, 5i** and **6i** calculated using the CB-Dock2 program.^[31]^ According to this analysis, numerous residues participate in hydrophobic interactions with the substrates, with only a limited contribution from specific hydrogen bonds (red dotted lines) (Vina score: < –7.0 kcal·mol^-1^). In addition, the overall shape of the active site is complementary to that of the ligands. Residues directly responsible for recognizing a docked intermediate are indicated in gray and those interacting only with other residues are displayed in green. Substrates **2i, 5i**, and **6i** are shown in magenta, orange, and cyan, respectively. **D**) Superposition of the three chains showing that the nucleophilic Ser83 is located at a suitable distance (2.9–3.0 Å) (yellow dotted lines in all panels) and angle (83° (**2**), 81° (**5**), and 71° (**6**))^[32]^ from the reactive carbonyl function of each compound.

When we attempted to dock SNAC thioesters **12i** and **14i** (Fig. S8), the CB-Dock2 calculation failed due to the high number of possible side-chain conformers. To surmount this problem, we truncated the lipidic side chains of both potential substrates to acetyl (which also yielded molecular volumes compatible in principle with that of the active site), and repeated the docking, alongside that of **13i**. All three yielded defined docking poses but the results were mediocre, due in the cases of **12i** and **14i** to a substantial number of steric clashes with cavity residues. Together, these data argue that a higher degree of branching adjacent to the dehydro-Ala impedes correct positioning of the chains within the active site.

In the case of **8i**, it differs from **6i** only by the presence of a longer aliphatic side chain. While it is accepted by the TE *in vivo*, it yielded poor docking results, presumably due to its overall size which exceeds that of the cavity volume. Our interpretation in this case is that, lacking branching, **8i** can be correctly positioned, but that the acyl portion must protrude beyond the active site. Taken together, these docking results argue that the reactivity trends of the engineered NRPSs as observed by NRPS engineering success reflect the specificity profile of the PxaA TE. We anticipate that in future, such limitations could be addressed by artificial intelligence-guided protein engineering of the TE active site.

## Conclusions

We have achieved the structural diversification of bacterial PAs and their indolizidine precursors by NRPS engineering. The full set of target derivatives was not obtained, however, likely due to the narrow substrate specificity of the TE domain, which prefers dipeptides bearing short-to medium-length acyl chains. Indeed, no PA derivatives with longer peptide chains have been identified to date in nature, and correspondingly, no PA-producing BGCs with more than two NRPS modules have been described in microbial genomes. Instead, natural variability in this metabolite family derives exclusively from selection of Ser or Thr as the first building block and the type of appended acyl chain, which originates from the fatty acid pool or is generated *de novo* by polyketide synthases (PKS) as in the case of some PA analogs from *Streptomyces*.^[14]^ This close structural similarity among PAs produced by species of both Gram-positive and Gram-negative bacteria, including pathogenic *Pseudomonas aeruginosa*, suggest a highly conserved ecological function. However, while several bioactivities for microbial PAs have been described in the literature,^[35– 37]^ a truly conserved target for this widespread NP class awaits identification. The reactivity of PAs with aldehydes as observed in the formation of bohemamine and pyrrolizwiline, and with nucleophiles in the case of functionalized PAs such as bohemamine, hints at the existence of several targets and thus potentially distinct modes of action.

## Author contributions

J.E. and S.K. performed all experiments, except for the modelling and docking experiments performed by A.C. K.J.W. and H.B.B. supervised the work, and J.E., A.C., K.J.W. and H.B.B. wrote the paper.

## Acknowledgements

Work in the Bode lab was supported by the Max Planck Society. Research in the S2BEAM lab was funded by ANR grant BiosynKADH (ANR-24-CE92-0032-02).

## Data availability statement

All data and materials can be found within the manuscript or in the supporting information.

