## Supplementary Methods, Tables & Figures for "Derivatization of the non-ribosomal peptide pyrrolizixenamide using NRPS engineering"

**Content**

**1. Material and Methods** **2**

**2. Supplementary Tables** **4**

**3. Supplementary Figures** **26**

**4. References** **34**

### 1. Material and Methods

#### Cultivation of strains

All *E. coli* and *Xenorhabdus* strains were cultivated in Lysogeny Broth (LB) medium (10 g/L tryptone, 5 g/L yeast extract, and 5 g/L NaCl) or in XPP3 medium[1] at 28 °C for *Xenorhabdus* and 37 °C for *E. coli*, with shaking at 200 rpm overnight. For cultivation on LB agar plates, 1.5% agar was added to the medium. Antibiotics (kanamycin (Km) [50 µg/mL], chloramphenicol (Cm) [34 µg/mL] and/or spectinomycin (Spec) [100 µg/mL] were added when appropriate. For long term conservation, 900 µL of the overnight culture were mixed with 600 µL of 50% (v/v) glycerol, then stored at −80 °C.

#### Plasmid assembly

Genomic DNA (gDNA) was purified from overnight (ON) cultures using the Monarch® Genomic DNA Purification Kit (New England Biolabs), according to the manufacturer's protocols. Plasmids were purified from *E. coli* ON cultures using the Monarch® Plasmid Miniprep Kit (New England Biolabs), according to the manufacturer's protocols. Polymerase chain reaction (PCR) was used to amplify DNA. Q5® High Fidelity DNA Polymerase or Phusion® Hot Start Flex DNA Polymerase, both from New England Biolabs, were used according to the manufacturer's instructions. PCR was conducted using a ProFlex™ 3 × 32-Well PCR System thermal cycler. For PCR amplification of full-length plasmid backbone, a subsequent digest was performed with *Dpn*I (Thermo Fisher Scientific). The PCR products were then subjected to electrophoresis at 120 V for 30 min on 1% TAE agarose gels. DNA fragments were separated using 1% TAE agarose gel electrophoresis at 120 V for 30 min. The fragments were then purified using the Monarch® DNA Gel Extraction Kit (New England Biolabs) according to the manufacturer's instructions. The NEBuilder® HiFi DNA Assembly Mix (New England Biolabs) was used for plasmid assembly according to the manufacturer's instructions. After transforming the assembly mix into *E. coli* DH10B::*mtaA*, the plasmids were verified by restriction digest using the appropriate restriction enzymes and the respective fragments were isolated and sequenced (MicroSynth). *E. coli* cells were transformed via electroporation.[18] For this, 1 mm cuvettes and a GenePulser® II (Bio-Rad Laboratories GmbH) set to 1250 V, 25 µF, and 200 Ω, were used. After electroporation, 500 µL of LB medium was added, and the cells were incubated at 37 °C with shaking for 1 h. Lastly, the cells were plated on LB agar plates containing the appropriate antibiotics and incubated overnight.

**Sample preparation for LC-MS analysis**

*E. coli* DH10B::*mtaA* production cultures were grown in XPP3[1] medium supplemented with 2% (v/v) XAD-16.[2] They were then incubated with shaking at 22 °C for 72 h. Metabolite production was induced at the beginning of cultivation using 0.2% L-arabinose. XAD-16 beads were harvested and incubated with one culture volume of methanol for 60 min at 180 rpm. Next, 500 µL of the MeOH phase was centrifuged for 20 min at 21,000 x *g*, and 100 µL of the supernatant were transferred to an HPLC vial for LC-MS analysis.

**LC-MS analysis**

Routine high-performance liquid chromatography-mass spectrometry (HPLC-MS) analysis was conducted on a Dionex Ultimate 3000 (Thermo Fisher Scientific) reversed-phase (RP)-HPLC system coupled to an AmaZonX (Bruker) electrospray ionization (ESI)-ion trap (IT) mass spectrometer. 5 µL samples were injected onto a C18 Acquity UPLC BEH column (Waters) and subjected to a 16 min gradient of 5–95% ACN/0.1% formic acid in $H_2O$/0.1% formic acid at a flow rate of 0.4 mL/min. The mass spectrometer was set to positive ionization mode with a mass range of 100–1200 *m*/*z* and a cone voltage of 4500 V. Further HPLC analysis was carried out using an Elute LC Series 1300 (Bruker) equipped with a C18 column (ACQUITY UPLC BEH C18, 130 Å, 2.1 mm × 50 mm, Waters). The separation was conducted at a flow rate of 0.4 mL/min using a 5–95% acetonitrile (ACN) gradient in water containing 0.1% formic acid (v/v) over 18 min. High-resolution mass spectrometric analyses were then performed using an ESI ion-trap mass spectrometer (timsTOF fleX MALDI 2, Bruker). ESI-MS spectra were recorded in positive ion mode with a mass range of *m*/*z* 100–2000. Data were analyzed using Data Analysis 4.3 and 6.1 software (Bruker). Unless otherwise stated, HPLC-MS chromatograms with the same X-axis have the same signal intensity.

***In silico* modeling of the PxaA TE domain and enzyme–substrate complexes**

Structural models of the native PxaA TE domain were generated using AlphaFold2[3] Peptide intermediates (*N*-acetylcysteamine derivatives) **2a**, **3a**, **6**, and **8**, as well as the metabolites derived from NRPS-44/45, -57/58, or -61/62, were docked into the active site using the CB-Dock2 program.[4] All figures were prepared using PyMOL (Schrödinger L.L.C. (2015 The PyMOL Molecular Graphics System, Version 1.8)).

### 2. Supplementary Tables

**Table S1.** Calculated and measured $[M+H]^+$ ions of compounds investigated in this work.

| Compound | Sum formula | $[M+H]^+$ ion obs'd., *m/z* | $[M+H]^+$ ion calc'd, *m/z* | Δppm |
|---|---|---|---|---|
| **1** | $C_{13}H_{20}N_2O_2$ | 237.1603 | 237.1597 | 2.4 |
| **2** | $C_{14}H_{20}N_2O_3$ | 265.1546 | 265.1547 | −0.3 |
| **5** | $C_{10}H_{12}N_2O_3$ | 209.0922 | 209.0920 | 0.7 |
| **6** | $C_{15}H_{22}N_2O_3$ | 279.1698 | 279.1703 | −0.7 |
| **7** | $C_{14}H_{22}N_2O_2$ | 251.1754 | 251.1754 | 0 |
| **8** | $C_{23}H_{38}N_2O_3$ | 391.2953 | 391.2955 | −0.6 |
| **9** | $C_{22}H_{38}N_2O_2$ | 363.3002 | 363.3006 | −1.2 |
| **10** | $C_{14}H_{22}N_2O_4$ | 283.1652 | 283.1652 | 0 |
| **11** | $C_{14}H_{22}N_2O_3$ | 267.1703 | 267.1703 | 0 |

**Table S2.** Strains used in this work.

| NRPS | Strain | | Genotype/Description | Reference |
|---|---|---|---|---|
| | | *X. stockiae* WT | *X. stockiae* DSM 17904 | DSMZ |
| | | *E. coli* DH10B | F–*mcr*A, Δ(*mrr*–*hsd*RMS-*mcr*BC) Φ80*lac*ZΔM15, Δ*lac*X74, *rec*A1, *end*A1, *ara*D139, Δ(*araleu*)7697 *gal*U, *gal*K, *rps*L, *nup*G, λ– | Invitrogen |
| | | *E. coli* DH10B::*mtaA* | *E. coli* DH10B with Δ*entD*::*mtaA* | [5] |
| | 10010 | *E. coli* DH10B::*mtaA* + pACYC_$P_{BAD}$_*pxaAB* | *E. coli* DH10B with Δ*entD*::*mtaA* + pACYC_$P_{BAD}$_*pxaAB* | [1] |
| XhpA | 10004 | *E. coli* DH10B::*mtaA* + *pACYC*_$P_{BAD}$_*xhpA* | *E. coli* DH10B with Δ*entD*::*mtaA* + pACYC_$P_{BAD}$_*xhpA* | [1] |
| PxaA | 10009 | *E. coli* DH10B::*mtaA* pACYC_ $P_{BAD}$_*pxaA* | *E. coli* DH10B with Δ*entD*::*mtaA* + pACYC_ $P_{BAD}$_*pxaA* | This work |
| | 10011 | *E. coli* DH10B::*mtaA* pCOLA_ $P_{BAD}$_*pxaB* | *E. coli* DH10B with Δ*entD*::*mtaA* + pACYC_ $P_{BAD}$_*pxaB* | This work |
| NRPS-64 | 10024 | *E. coli* DH10B::*mtaA* pACYC_ $P_{BAD}$_*pxaA*_R | *E. coli* DH10B with Δ*entD*::*mtaA* + pACYC_ $P_{BAD}$_*pxaA*_R | This work |
| NRPS-65 | 10025 | *E. coli* DH10B::*mtaA* pACYC_ $P_{BAD}$_*pxaA*_T2stop | *E. coli* DH10B with Δ*entD*::*mtaA* + pACYC_ $P_{BAD}$_*pxaA*_T2stop | This work |
| NRPS-66 | 10059 | *E. coli* DH10B::*mtaA* pACYC_ $P_{BAD}$_*pxaA*_TE-S83A | *E. coli* DH10B with Δ*entD*::*mtaA* + pACYC_ $P_{BAD}$_*pxaA*_TE-S83A | This work |
| NRPS-7 | 10060 | *E. coli* DH10B::*mtaA* pJE6 | *E. coli* DH10B with Δ*entD*::*mtaA* + pJE6 | This work |
| NRPS-13 | 10061 | *E. coli* DH10B::*mtaA* pJE8 | *E. coli* DH10B with Δ*entD*::*mtaA* + pJE8 | This work |
| NRPS-1 | 10062 | *E. coli* DH10B::*mtaA* pJE2 | *E. coli* DH10B with Δ*entD*::*mtaA* + pJE2 | This work |
| NRPS-3 | 10063 | *E. coli* DH10B::*mtaA* pJE4 | *E. coli* DH10B with Δ*entD*::*mtaA* + pJE4 | This work |
| NRPS-6 | 10064 | *E. coli* DH10B::*mtaA* pJE5 | *E. coli* DH10B with Δ*entD*::*mtaA* + pJE5 | This work |

| | | | | |
|---|---|---|---|---|
| NRPS-11 | 10066 | *E. coli* DH10B::*mtaA* pJE1 | *E. coli* DH10B with Δ*entD*::*mtaA* + pJE1 | This work |
| NRPS-2 | 1006 | *E. coli* DH10B::*mtaA* pJE3 | *E. coli* DH10B with Δ*entD*::*mtaA* + pJE3 | This work |
| NRPS-14 | 10068 | *E. coli* DH10B::*mtaA* pJE7 | *E. coli* DH10B with Δ*entD*::*mtaA* + pJE7 | This work |
| NRPS-8 | 10071 | *E. coli* DH10B::*mtaA* pJE14 | *E. coli* DH10B with Δ*entD*::*mtaA* + pJE14 | This work |
| NRPS-12 | 10100 | *E. coli* DH10B::*mtaA* pJE13 | *E. coli* DH10B with Δ*entD*::*mtaA* + pJE13 | This work |
| NRPS-4 | 10106 | *E. coli* DH10B::*mtaA* pJE39 | *E. coli* DH10B with Δ*entD*::*mtaA* + pJE39 | This work |
| NRPS-5 | 10107 | *E. coli* DH10B::*mtaA* pJE40 | *E. coli* DH10B with Δ*entD*::*mtaA* + pJE40 | This work |
| NRPS-9 | 10108 | *E. coli* DH10B::*mtaA* pJE41 | *E. coli* DH10B with Δ*entD*::*mtaA* + pJE41 | This work |
| NRPS-10 | 10109 | *E. coli* DH10B::*mtaA* pJE42 | *E. coli* DH10B with Δ*entD*::*mtaA* + pJE42 | This work |
| NRPS-15 Part 1 | 10132 | *E. coli* DH10B::*mtaA* pJE43 | *E. coli* DH10B with Δ*entD*::*mtaA* + pJE43 | This work |
| NRPS-17 Part 1 | 10133 | *E. coli* DH10B::*mtaA* pJE44 | *E. coli* DH10B with Δ*entD*::*mtaA* + pJE44 | This work |
| NRPS-17 Part 2 | 10134 | *E. coli* DH10B::*mtaA* pJE45 | *E. coli* DH10B with Δ*entD*::*mtaA* + pJE45 | This work |
| | 10154 | *E. coli* DH10B::*mtaA* pJE56 + pPC1018 | *E. coli* DH10B with Δ*entD*::*mtaA* + pJE56 + pPC1018 | This work, [6] |
| | 10155 | *E. coli* DH10B::*mtaA* pJE57 + pPC1017 | *E. coli* DH10B with Δ*entD*::*mtaA* + pJE57 + pPC1017 | This work, [6] |
| NRPA-16 Part 1 | 10170 | *E. coli* DH10B::*mtaA* pJE58 | *E. coli* DH10B with Δ*entD*::*mtaA* + pJE58 | This work |
| NRPS-18 Part 1 | 10171 | *E. coli* DH10B::*mtaA* pJE59 | *E. coli* DH10B with Δ*entD*::*mtaA* + pJE59 | This work |
| NRPS-18 Part 2 | 10173 | *E. coli* DH10B::*mtaA* pJE61 | *E. coli* DH10B with Δ*entD*::*mtaA* + pJE61 | This work |
| NRPS-32 | 10102 | *E. coli* DH10B::*mtaA* pJE27 | *E. coli* DH10B with Δ*entD*::*mtaA* + pJE27 | This work |

| | | | | |
|---|---|---|---|---|
| NRPS-31 | 10103 | *E. coli* DH10B::*mtaA* pJE28 | *E. coli* DH10B with Δ*entD*::*mtaA* + pJE28 | This work |
| NRPS-30 | 10104 | *E. coli* DH10B::*mtaA* pJE29 | *E. coli* DH10B with Δ*entD*::*mtaA* + pJE29 | This work |
| NRPS-33 | 10105 | *E. coli* DH10B::*mtaA* pJE30 | *E. coli* DH10B with Δ*entD*::*mtaA* + pJE30 | This work |
| NRPS-64 | 10354 | *E. coli* DH10B::*mtaA* pJE109 | *E. coli* DH10B with Δ*entD*::*mtaA* + pJE109 | This work |
| NRPS-63 | 10355 | *E. coli* DH10B::*mtaA* pJE110 | *E. coli* DH10B with Δ*entD*::*mtaA* + pJE110 | This work |
| NRPS-38 | 10138 | *E. coli* DH10B::*mtaA* pJE47 | *E. coli* DH10B with Δ*entD*::*mtaA* + pJE47 | This work |
| NRPS-39 | 10139 | *E. coli* DH10B::*mtaA* pJE48 | *E. coli* DH10B with Δ*entD*::*mtaA* + pJE48 | This work |
| NRPS-40 | 10136 | *E. coli* DH10B::*mtaA* pJE49 | *E. coli* DH10B with Δ*entD*::*mtaA* + pJE49 | This work |
| NRPS-41 | 1014 | *E. coli* DH10B::*mtaA* pJE50 | *E. coli* DH10B with Δ*entD*::*mtaA* + pJE50 | This work |
| NRPS-42 | 10137 | *E. coli* DH10B::*mtaA* pJE51 | *E. coli* DH10B with Δ*entD*::*mtaA* + pJE51 | This work |
| NRPS-43 | 10140 | *E. coli* DH10B::*mtaA* pJE52 | *E. coli* DH10B with Δ*entD*::*mtaA* + pJE52 | This work |
| NRPS-44 | 10141 | *E. coli* DH10B::*mtaA* pJE53 | *E. coli* DH10B with Δ*entD*::*mtaA* + pJE53 | This work |
| NRPS-45 | 10142 | *E. coli* DH10B::*mtaA* pJE54 | *E. coli* DH10B with Δ*entD*::*mtaA* + pJE54 | This work |
| NRPS-46 | 10143 | *E. coli* DH10B::*mtaA* pJE55 | *E. coli* DH10B with Δ*entD*::*mtaA* + pJE55 | This work |
| NRPS-47 | 10147 | *E. coli* DH10B::*mtaA* pJE56 | *E. coli* DH10B with Δ*entD*::*mtaA* + pJE56 | This work |
| NRPS-48 | 10148 | *E. coli* DH10B::*mtaA* pJE57 | *E. coli* DH10B with Δ*entD*::*mtaA* + pJE57 | This work |
| | 10086 | *E. coli* DH10B::*mtaA* pJE_frs5 | *E. coli* DH10B with Δ*entD*::*mtaA* + pJE_frs5 | This work |
| | 10089 | *E. coli* DH10B::*mtaA* pJE_frs11 | *E. coli* DH10B with Δ*entD*::*mtaA* + pJE_frs11 | This work |
| NRPS-36 | 10076 | *E. coli* DH10B::*mtaA* pJE23 | *E. coli* DH10B with Δ*entD*::*mtaA* + pJE23 | This work |
| NRPS-37 | 10077 | *E. coli* DH10B::*mtaA* pJE24 | *E. coli* DH10B with Δ*entD*::*mtaA* + pJE24 | This work |
| NRPS-34 | 10078 | *E. coli* DH10B::*mtaA* pJE25 | *E. coli* DH10B with Δ*entD*::*mtaA* + pJE25 | This work |
| NRPS-35 | 10079 | *E. coli* DH10B::*mtaA* pJE26 | *E. coli* DH10B with Δ*entD*::*mtaA* + pJE26 | This work |
| NRPS-54 | 10297 | *E. coli* DH10B::*mtaA* pJE76 | *E. coli* DH10B with Δ*entD*::*mtaA* + pJE76 | This work |
| NRPS-52 | 10298 | *E. coli* DH10B::*mtaA* pJE77 | *E. coli* DH10B with Δ*entD*::*mtaA* + pJE77 | This work |
| NRPS-53 | 10300 | *E. coli* DH10B::*mtaA* pJE80 | *E. coli* DH10B with Δ*entD*::*mtaA* + pJE80 | This work |

| | | | | |
|---|---|---|---|---|
| NRPS-49 | 10304 | *E. coli* DH10B::*mtaA* pJE79 | *E. coli* DH10B with Δ*entD*::*mtaA* + pJE79 | This work |
| NRPS-51 | 10301 | *E. coli* DH10B::*mtaA* pJE81 | *E. coli* DH10B with Δ*entD*::*mtaA* + pJE81 | This work |
| NRPS-55 | 10305 | *E. coli* DH10B::*mtaA* pJE86 | *E. coli* DH10B with Δ*entD*::*mtaA* + pJE86 | This work |
| NRPS-58 | 10306 | *E. coli* DH10B::*mtaA* pJE87 | *E. coli* DH10B with Δ*entD*::*mtaA* + pJE87 | This work |
| NRPS-57 | 10307 | *E. coli* DH10B::*mtaA* pJE88 | *E. coli* DH10B with Δ*entD*::*mtaA* + pJE88 | This work |
| NRPS-62 | 10308 | *E. coli* DH10B::*mtaA* pJE89 | *E. coli* DH10B with Δ*entD*::*mtaA* + pJE89 | This work |
| NRPS-60 | 10309 | *E. coli* DH10B::*mtaA* pJE91 | *E. coli* DH10B with Δ*entD*::*mtaA* + pJE91 | This work |
| NRPS-59 | 10310 | *E. coli* DH10B::*mtaA* pJE92 | *E. coli* DH10B with Δ*entD*::*mtaA* + pJE92 | This work |
| NRPS-19 | 10174 | *E. coli* DH10B::*mtaA* pJE45 + pJW77 | *E. coli* DH10B with Δ*entD*::*mtaA* + pJE45 + pJW77 | This work, [7] |
| NRPS-20 | 1017 | *E. coli* DH10B::*mtaA* pJE45 + pJW91 | *E. coli* DH10B with Δ*entD*::*mtaA* + pJE45 + pJW91 | This work, [7] |
| NRPS-22 | 10176 | *E. coli* DH10B::*mtaA* pJE45 + pJW92 | *E. coli* DH10B with Δ*entD*::*mtaA* + pJE45 + pJW92 | This work, [7] |
| NRPS-23 | 10177 | *E. coli* DH10B::*mtaA* pJE45 + pJW93 | *E. coli* DH10B with Δ*entD*::*mtaA* + pJE45 + pJW93 | This work, [7] |
| NRPS-24 | 10178 | *E. coli* DH10B::*mtaA* pJE45 + pJW114 | *E. coli* DH10B with Δ*entD*::*mtaA* + pJE45 + pJW114 | This work, [7] |
| NRPS-21 | 10181 | *E. coli* DH10B::*mtaA* pJE45 + pNA128 | *E. coli* DH10B with Δ*entD*::*mtaA* + pJE45 + pNA128 | This work, [7] |
| NRPS-25 Part 1 | 10196 | *E. coli* DH10B::*mtaA* pJE64 | *E. coli* DH10B with Δ*entD*::*mtaA* + pJE64 | This work |
| NRPS-26 Part 1 | 10197 | *E. coli* DH10B::*mtaA* pJE65 | *E. coli* DH10B with Δ*entD*::*mtaA* + pJE65 | This work |
| NRPS-27 Part 1 | 10198 | *E. coli* DH10B::*mtaA* pJE66 | *E. coli* DH10B with Δ*entD*::*mtaA* + pJE66 | This work |
| NRPS-28 Part 1 | 10199 | *E. coli* DH10B::*mtaA* pJE67 | *E. coli* DH10B with Δ*entD*::*mtaA* + pJE67 | This work |
| NRPS-29 Part 1 | 10200 | *E. coli* DH10B::*mtaA* pJE68 | *E. coli* DH10B with Δ*entD*::*mtaA* + pJE68 | This work |
| NRPS-65 | 10349 | *E. coli* DH10B::*mtaA* pJE103 | *E. coli* DH10B with Δ*entD*::*mtaA* + pJE103 | This work |

**Table S3.** Plasmids used in this work.

| Plasmid | Description | Reference |
|---|---|---|
| pCK_0431 pACYC | *ori* p15A, cm$^R$, *araC*, $P_{BAD}$ and *tacI*, I-*Sce*I, I-*Ceu*I, | C. Kegler, [1] |
| pCK_0433 pCOLA | *ori* ColA, kan$^R$, araC, $P_{BAD}$, *tacI*, I-*Sce*I, I-*Ceu*I, minus *Bsa*I | C. Kegler, [1] |
| pACYC_$P_{BAD}$_*pxaAB* | *ori* p15A, cm$^R$, *araC*, $P_{BAD}$ and *tacI*, I-*Sce*I, I-*Ceu*I, *pxaAB* inserted under $P_{BAD}$ control | This work |
| pACYC_$P_{BAD}$_*xhpA* | *ori* p15A, cm$^R$, *araC*, $P_{BAD}$ and *tacI*, I-*Sce*I, I-*Ceu*I, *xhpA* inserted under $P_{BAD}$ control | This work |
| pACYC_ $P_{BAD}$_*pxaA* | *ori* p15A, cm$^R$, *araC*, $P_{BAD}$ and *tacI*, I-*Sce*I, I-*Ceu*I, *pxaA* inserted under $P_{BAD}$ control | This work |
| pACYC_ $P_{BAD}$_*pxaA*_R | *ori* p15A, cm$^R$, *araC*, $P_{BAD}$ and *tacI*, I-*Sce*I, I-*Ceu*I, *pxaA*-$C_1$–$T_2$_*xtvB*-R inserted under $P_{BAD}$ control | This work |
| pACYC_ $P_{BAD}$_*pxaA*_T2stop | *ori* p15A, cm$^R$, *araC*, $P_{BAD}$ and *tacI*, I-*Sce*I, I-*Ceu*I, *pxaA*-$C_1$–$T_2$ inserted under $P_{BAD}$ control | This work |
| pACYC_ $P_{BAD}$_*pxaA*_TE-S83A | *ori* p15A, cm$^R$, *araC*, $P_{BAD}$ and *tacI*, I-*Sce*I, I-*Ceu*I, *pxaA*_ TE-S83A inserted under $P_{BAD}$ control | This work |
| pJE6 | *ori* p15A, cm$^R$, *araC*, $P_{BAD}$ and *tacI*, I-*Sce*I, I-*Ceu*I, *xhpA*-$C_1$–$A_1$_*pxaA*-$T_1$–TE inserted under $P_{BAD}$ control | This work |
| pJE8 | *ori* p15A, cm$^R$, *araC*, $P_{BAD}$ and *tacI*, I-*Sce*I, I-*Ceu*I, *xhpA*-$C_1$–$T_1$(XUT$^{IV}$)_*pxaA*-$T_1$(XUT$^{IV}$)–TE inserted under $P_{BAD}$ control | This work |
| pJE2 | *ori* p15A, cm$^R$, *araC*, $P_{BAD}$ and *tacI*, I-*Sce*I, I-*Ceu*I, *pxaA*-$C_1$–$T_1$(XUT$^{I}$)_*xhpA*-$T_1$(XUT$^{I}$)–TE inserted under $P_{BAD}$ control | This work |
| pJE4 | *ori* p15A, cm$^R$, *araC*, $P_{BAD}$ and *tacI*, I-*Sce*I, I-*Ceu*I, *pxaA*-$C_1$–$T_1$(XUT$^{IV}$)_*xhpA*-$T_1$(XUT$^{IV}$)–TE inserted under $P_{BAD}$ control | This work |
| pJE5 | *ori* p15A, cm$^R$, *araC*, $P_{BAD}$ and *tacI*, I-*Sce*I, I-*Ceu*I, *xhpA*-$C_1$_*pxaA*-$A_1$–TE inserted under $P_{BAD}$ control | This work |
| pJE1 | *ori* p15A, cm$^R$, *araC*, $P_{BAD}$ and *tacI*, I-*Sce*I, I-*Ceu*I, *pxaA*-$C_1$_*xhpA*-$A_1$–TE inserted under $P_{BAD}$ control | This work |
| pJE3 | *ori* p15A, cm$^R$, *araC*, $P_{BAD}$ and *tacI*, I-*Sce*I, I-*Ceu*I, *pxaA*-$C_1$–$T_1$(XUT$^{III}$)_*xhpA*-$T_1$(XUT$^{III}$)–TE inserted under $P_{BAD}$ control | This work |
| pJE7 | *ori* p15A, cm$^R$, *araC*, $P_{BAD}$ and *tacI*, I-*Sce*I, I-*Ceu*I, *xhpA*-$C_1$–$T_1$(XUT$^{III}$)_*pxaA*-$T_1$(XUT$^{III}$)–TE inserted under $P_{BAD}$ control | This work |
| pJE14 | *ori* p15A, cm$^R$, *araC*, $P_{BAD}$ and *tacI*, I-*Sce*I, I-*Ceu*I, *xhpA*-$C_1$–$C_2$_*pxaA*-$A_2$–TE inserted under $P_{BAD}$ control | This work |
| pJE13 | *ori* p15A, cm$^R$, *araC*, $P_{BAD}$ and *tacI*, I-*Sce*I, I-*Ceu*I, *pxaA*-$C_1$–$C_2$_*xhpA*-$A_2$–TE inserted under $P_{BAD}$ control | This work |

| | | |
|---|---|---|
| pJE39 | *ori* p15A, $cm^R$, *araC*, $P_{BAD}$ and *tacI*, I-*Sce*I, I-*Ceu*I, *pxaA*-$C_1$–$A_2$_*xhpA*-$T_2$–TE inserted under $P_{BAD}$ control | This work |
| pJE40 | *ori* p15A, $cm^R$, *araC*, $P_{BAD}$ and *tacI*, I-*Sce*I, I-*Ceu*I, *pxaA*-$C_1$–$T_2$($XUT^{III}$)_*xhpA*-$T_2$($XUT^{III}$)_–TE inserted under $P_{BAD}$ control | This work |
| pJE41 | *ori* p15A, $cm^R$, *araC*, $P_{BAD}$ and *tacI*, I-*Sce*I, I-*Ceu*I, *xhpA*-$C_1$–$A_2$_*pxaA*-$T_2$–TE inserted under $P_{BAD}$ control | This work |
| pJE42 | *ori* p15A, $cm^R$, *araC*, $P_{BAD}$ and *tacI*, I-*Sce*I, I-*Ceu*I, *xhpA*-$C_1$–$T_2$($XUT^{III}$)_*pxaA*-$T_2$($XUT^{III}$)_–TE inserted under $P_{BAD}$ control | This work |
| pJE22 | *ori* ColA, $kan^R$, araC, $P_{BAD}$, *tacI*, I-*Sce*I, I-*Ceu*I, minus *Bsa*I, SZ18_*pxaA*-$T_1$–TE inserted under $P_{BAD}$ control | This work |
| pJE43 | *ori* p15A, $cm^R$, *araC*, $P_{BAD}$ and *tacI*, I-*Sce*I, I-*Ceu*I, *pxaA*-$C_1$_SZ17 inserted under $P_{BAD}$ control | This work |
| pJE44 | *ori* p15A, $cm^R$, *araC*, $P_{BAD}$ and *tacI*, I-*Sce*I, I-*Ceu*I, *pxaA*-$C_1$–$C_2$_SZ17 inserted under $P_{BAD}$ control | This work |
| pJE45 | *ori* ColA, $kan^R$, araC, $P_{BAD}$, *tacI*, I-*Sce*I, I-*Ceu*I, minus *Bsa*I, SZ18_*pxaA*-$A_1$–TE inserted under $P_{BAD}$ control | This work |
| pJE46 | *ori* ColA, $kan^R$, araC, $P_{BAD}$, *tacI*, I-*Sce*I, I-*Ceu*I, minus *Bsa*I, SZ18_*xhpA*-$A_2$–TE inserted under $P_{BAD}$ control | This work |
| pPC1018 | *ori ColA, $kan^R$, araC-* $P_{BAD}$ *odlB and tacI* | [6] |
| pPC1017 | *ori ColA, $kan^R$, araC-* $P_{BAD}$ *odlF and tacI* | [6] |
| pJE58 | *ori* p15A, $cm^R$, *araC*, $P_{BAD}$ and *tacI*, I-*Sce*I, I-*Ceu*I, *pxaA*-$C_1$–$A_1$_SZ17 inserted under $P_{BAD}$ control | This work |
| pJE59 | *ori* p15A, $cm^R$, *araC*, $P_{BAD}$ and *tacI*, I-*Sce*I, I-*Ceu*I, *pxaA*-$C_1$–$A_2$_SZ17 inserted under $P_{BAD}$ control | This work |
| pJE61 | *ori* ColA, $kan^R$, araC, $P_{BAD}$, *tacI*, I-*Sce*I, I-*Ceu*I, minus *Bsa*I, SZ18_*pxaA*-$T_2$-TE inserted under $P_{BAD}$ control | This work |
| pJE27 | *ori* p15A, $cm^R$, *araC*, $P_{BAD}$ and *tacI*, I-*Sce*I, I-*Ceu*I, *txlA*-$C_1$ _*pxaA*-$TA_1$–TE inserted under $P_{BAD}$ control | This work |
| pJE28 | *ori* p15A, $cm^R$, *araC*, $P_{BAD}$ and *tacI*, I-*Sce*I, I-*Ceu*I, *txlA*-$C_1$–$A_1$ _*pxaA*-$T_1$–TE inserted under $P_{BAD}$ control | This work |
| pJE29 | *ori* p15A, $cm^R$, *araC*, $P_{BAD}$ and *tacI*, I-*Sce*I, I-*Ceu*I, *xabA*-$C_1$ _*pxaA*-$A_1$–TE inserted under $P_{BAD}$ control | This work |
| pJE30 | *ori* p15A, $cm^R$, *araC*, $P_{BAD}$ and *tacI*, I-*Sce*I, I-*Ceu*I, *xabA*-$C_1$–$A_1$_*pxaA*-$T_1$–TE inserted under $P_{BAD}$ control | This work |
| pJE109 | *ori* p15A, $cm^R$, *araC*, $P_{BAD}$ and *tacI*, I-*Sce*I, I-*Ceu*I, *xld*-C1_*txlA*-$A_1$_*pxaA*-$T_1$–TE inserted under $P_{BAD}$ control | This work |

| | | |
|---|---|---|
| pJE110 | *ori* p15A, cm$^{R}$, *araC*, $P_{BAD}$ and *tacI*, I-*Sce*I, I-*Ceu*I, *xld*-C1_*xabA*-A$_1$_*pxaA*-T$_1$–TE inserted under $P_{BAD}$ control | This work |
| pJE47 | *ori* p15A, cm$^{R}$, *araC*, $P_{BAD}$ and *tacI*, I-*Sce*I, I-*Ceu*I, *paxA*-C$_1$_*pxaA*-A$_1$–TE inserted under $P_{BAD}$ control | This work |
| pJE48 | *ori* p15A, cm$^{R}$, *araC*, $P_{BAD}$ and *tacI*, I-*Sce*I, I-*Ceu*I, *paxA*-C$_1$–A$_1$_*pxaA*-T$_1$–TE inserted under $P_{BAD}$ control | This work |
| pJE49 | *ori* p15A, cm$^{R}$, *araC*, $P_{BAD}$ and *tacI*, I-*Sce*I, I-*Ceu*I, *szeS*-C$_1$–C$_2$_*pxaA*-A$_1$–TE inserted under $P_{BAD}$ control | This work |
| pJE50 | *ori* p15A, cm$^{R}$, *araC*, $P_{BAD}$ and *tacI*, I-*Sce*I, I-*Ceu*I, *szeS*-C$_1$–A$_2$_*pxaA*-T$_1$–TE inserted under $P_{BAD}$ control | This work |
| pJE51 | *ori* p15A, cm$^{R}$, *araC*, $P_{BAD}$ and *tacI*, I-*Sce*I, I-*Ceu*I, *XinnSplit*-C$_1$–C$_2$_*pxaA*-A$_1$–TE inserted under $P_{BAD}$ control | This work |
| pJE52 | *ori* p15A, cm$^{R}$, *araC*, $P_{BAD}$ and *tacI*, I-*Sce*I, I-*Ceu*I, *XinnSplit*-C$_1$–A$_2$_*pxaA*-T$_1$–TE inserted under $P_{BAD}$ control | This work |
| pJE53 | *ori* p15A, cm$^{R}$, *araC*, $P_{BAD}$ and *tacI*, I-*Sce*I, I-*Ceu*I, *xld*-C$_1$–C$_3$_*pxaA*-A$_1$–TE inserted under $P_{BAD}$ control | This work |
| pJE54 | *ori* p15A, cm$^{R}$, *araC*, $P_{BAD}$ and *tacI*, I-*Sce*I, I-*Ceu*I, *xld*-C$_1$–A$_3$_*pxaA*-T$_1$–TE inserted under $P_{BAD}$ control | This work |
| pJE55 | *ori* p15A, cm$^{R}$, *araC*, $P_{BAD}$ and *tacI*, I-*Sce*I, I-*Ceu*I, *frsA*-C$_1$–A$_1$_*pxaA*-T$_1$–TE inserted under $P_{BAD}$ control | This work |
| pJE56 | *ori* p15A, cm$^{R}$, *araC*, $P_{BAD}$ and *tacI*, I-*Sce*I, I-*Ceu*I, *odlS1*-C$_1$–A$_2$_*pxaA*-T$_1$–TE inserted under $P_{BAD}$ control | This work |
| pJE57 | *ori* p15A, cm$^{R}$, *araC*, $P_{BAD}$ and *tacI*, I-*Sce*I, I-*Ceu*I, *odlS1*-C$_1$–A$_2$_*odlS4*-T$_1$–A$_1$_*pxaA*-T$_1$–TE inserted under $P_{BAD}$ control | This work |
| pJE_frs5 | CloDF13 ori, Spec$^{R}$, *araC*, $P_{BAD}$ and *tacI*, I-*Sce*I, I-*Ceu*I, T7-term, *frsH* inserted under $P_{BAD}$ control | This work |
| pJE_frs11 | *ori* ColA, kan$^{R}$, araC, $P_{BAD}$, *tacI*, I-*Sce*I, I-*Ceu*I, minus *Bsa*I, *frsB* inserted under $P_{BAD}$ control | This work |
| pJE23 | *ori* p15A, cm$^{R}$, *araC*, $P_{BAD}$ and *tacI*, I-*Sce*I, I-*Ceu*I, *ambS*-C$_1$_*pxaA*-A$_1$–TE inserted under $P_{BAD}$ control | This work |
| pJE24 | *ori* p15A, cm$^{R}$, *araC*, $P_{BAD}$ and *tacI*, I-*Sce*I, I-*Ceu*I, *ambS*-C$_1$–A$_1$_*pxaA*-T$_1$–TE inserted under $P_{BAD}$ control | This work |
| pJE25 | *ori* p15A, cm$^{R}$, *araC*, $P_{BAD}$ and *tacI*, I-*Sce*I, I-*Ceu*I, *ambS*-C$_1$_*pxaA*-A$_1$–TE inserted under $P_{BAD}$ control | This work |
| pJE26 | *ori* p15A, cm$^{R}$, *araC*, $P_{BAD}$ and *tacI*, I-*Sce*I, I-*Ceu*I, *ambS*-C$_1$–A$_1$_*pxaA*-T$_1$–TE inserted under $P_{BAD}$ control | This work |
| pJE76 | *ori* p15A, cm$^{R}$, *araC*, $P_{BAD}$ and *tacI*, I-*Sce*I, I-*Ceu*I, *pttB*-C$_1$_*txlA*-A$_1$_*pxaA*-T$_1$–TE inserted under $P_{BAD}$ control | This work |

| | | |
|---|---|---|
| pJE80 | *ori* p15A, cm$^R$, *araC*, $P_{BAD}$ and *tacI*, I-*Sce*I, I-*Ceu*I, *pttB*-$C_1$_*xabA*-$A_1$_*pxaA*-$T_1$–TE inserted under $P_{BAD}$ control | This work |
| pJE75 | *ori* p15A, cm$^R$, *araC*, $P_{BAD}$ and *tacI*, I-*Sce*I, I-*Ceu*I, *paxA*-$C_1$_*txlA*-$A_1$_*pxaA*-$T_1$–TE inserted under $P_{BAD}$ control | This work |
| pJE79 | *ori* p15A, cm$^R$, *araC*, $P_{BAD}$ and *tacI*, I-*Sce*I, I-*Ceu*I, *paxA*-$C_1$_*xabA*-$A_1$_*pxaA*-$T_1$–TE inserted under $P_{BAD}$ control | This work |
| pJE77 | *ori* p15A, cm$^R$, *araC*, $P_{BAD}$ and *tacI*, I-*Sce*I, I-*Ceu*I, *pxaA*-$C_1$_*txlA*-$A_1$_*pxaA*-$T_1$–TE inserted under $P_{BAD}$ control | This work |
| pJE81 | *ori* p15A, cm$^R$, *araC*, $P_{BAD}$ and *tacI*, I-*Sce*I, I-*Ceu*I, *pxaA*-$C_1$_*xabA*-$A_1$_*pxaA*-$T_1$–TE inserted under $P_{BAD}$ control | This work |
| pJE86 | *ori* p15A, cm$^R$, *araC*, $P_{BAD}$ and *tacI*, I-*Sce*I, I-*Ceu*I, *txlA*-$C_1$-$C_2$_*xabA*-$A_1$_*pxaA*-$T_1$–TE inserted under $P_{BAD}$ control | This work |
| pJE87 | *ori* p15A, cm$^R$, *araC*, $P_{BAD}$ and *tacI*, I-*Sce*I, I-*Ceu*I, *fitAB*-$C_1$-$C_2$_*txlA*-$A_1$_*pxaA*-$T_1$–TE inserted under $P_{BAD}$ control | This work |
| pJE88 | *ori* p15A, cm$^R$, *araC*, $P_{BAD}$ and *tacI*, I-*Sce*I, I-*Ceu*I, *fitAB*-$C_1$-$C_2$_*xabA*-$A_1$_*pxaA*-$T_1$–TE inserted under $P_{BAD}$ control | This work |
| pJE89 | *ori* p15A, cm$^R$, *araC*, $P_{BAD}$ and *tacI*, I-*Sce*I, I-*Ceu*I, *xldS*-$C_1$-$C_3$_*txlA*-$A_1$_*pxaA*-$T_1$–TE inserted under $P_{BAD}$ control | This work |
| pJE90 | *ori* p15A, cm$^R$, *araC*, $P_{BAD}$ and *tacI*, I-*Sce*I, I-*Ceu*I, *xldS*-$C_1$-$C_3$_*xabA*-$A_1$_*pxaA*-$T_1$–TE inserted under $P_{BAD}$ control | This work |
| pJE91 | *ori* p15A, cm$^R$, *araC*, $P_{BAD}$ and *tacI*, I-*Sce*I, I-*Ceu*I, *fitAB*-$C_1$-$C_3$_*txlA*-$A_1$_*pxaA*-$T_1$–TE inserted under $P_{BAD}$ control | This work |
| pJE92 | *ori* p15A, cm$^R$, *araC*, $P_{BAD}$ and *tacI*, I-*Sce*I, I-*Ceu*I, *fitAB*-$C_1$-$C_3$_*xabA*-$A_1$_*pxaA*-$T_1$–TE inserted under $P_{BAD}$ control | This work |
| pJW77 | *ori p15A, cm$^R$, araC-$P_{BAD}$ bicA*_$A_1T_1C/E_2A_2T_2C_3$-SZ17 and *tacI-araE* | |
| pJW91 | *ori p15A, cm$^R$, araC-$P_{BAD}$ ambS*_ $A_1T_1C/E_2A_2T_2C_3$-SZ17 and *tacI-araE* | |
| pJW92 | *ori p15A, cm$^R$, araC-$P_{BAD}$ szeS_FtA1T1C/E2A2T2C3*-SZ17 and *tacI-araE* | [8] |
| pJW93 | *ori p15A, cm$^R$, araC-$P_{BAD}$ xldS*_$C_1A_1T_1C/E_2A_2T_2C_3$-SZ17 and *tacI-araE* | |
| pJW114 | *ori p15A, cm$^R$, araC-$P_{BAD}$ bacA*_$A_1T_1CyA_2T_2C_3$- SZ17and *tacI-araE* | |
| pNA128 | ori p15A, cm$^R$, *araC-PBAD XldS*_C1-t1/2_*xabA*_T1/2-C2- SZ17 and *tac*I-*araE* | [7] |
| pJE64 | *ori* p15A, cm$^R$, *araC*, $P_{BAD}$ and *tacI*, I-*Sce*I, I-*Ceu*I, *paxA*-$C_1$_SZ17 inserted under $P_{BAD}$ control | This work |

| | | |
|---|---|---|
| pJE65 | *ori* p15A, cm$^R$, *araC*, $P_{BAD}$ and *tacI*, I-*Sce*I, I-*Ceu*I, *pttB*-$C_1$_SZ17 inserted under $P_{BAD}$ control | This work |
| pJE66 | *ori* p15A, cm$^R$, *araC*, $P_{BAD}$ and *tacI*, I-*Sce*I, I-*Ceu*I, *XinnSplit*-$C_1$_SZ17 inserted under $P_{BAD}$ control | This work |
| pJE67 | *ori* p15A, cm$^R$, *araC*, $P_{BAD}$ and *tacI*, I-*Sce*I, I-*Ceu*I, *SzeS*-$C_1$_SZ17 inserted under $P_{BAD}$ control | This work |
| pJE68 | *ori* p15A, cm$^R$, *araC*, $P_{BAD}$ and *tacI*, I-*Sce*I, I-*Ceu*I, *xldS*-$C_1$_SZ17 inserted under $P_{BAD}$ control | This work |
| pJE103 | *ori* p15A, cm$^R$, *araC*, $P_{BAD}$ and *tacI*, I-*Sce*I, I-*Ceu*I, *pxaA*-$C_1$–$T_{2half}$_*gxpS*-$T_{half}$-*TE* inserted under $P_{BAD}$ control | This work |
| pJE104 | *ori* p15A, cm$^R$, *araC*, $P_{BAD}$ and *tacI*, I-*Sce*I, I-*Ceu*I, *pxaA*-$C_1$–$A_2$_*gxpS-T-TE* inserted under $P_{BAD}$ control | This work |

**Table S4.** Oligonucleotides used in this work.

| **Plasmid** | **Oligonucleotide** | **Sequence 5′ → 3′** | **PCR template** |
|---|---|---|---|
| pACYC_$P_{BAD}$_*pxaAB* | JE_G_for3.1 | GTTTTTTTGGGCTAACAGGAGGAATTCCATGAAAACTTCACAATTAGTACCTCTTACC | *X. stockiae* gDNA |
| | JE_G_rev6.1 | GCAGCAGCCTAGGTTAATTAATTATAGGGTCTGGAGAGTAAGATCTC | |
| pACYC_ $P_{BAD}$_*pxaA* | JE_G_for3.1 | GTTTTTTTGGGCTAACAGGAGGAATTCCATGAAAACTTCACAATTAGTACCTCTTACC | *X. stockiae* gDNA |
| | JE_G_rev5.1 | GTTTTTTTGGGCTAACAGGAGGAATTCCATGAGTAAAAATCCAACTGCGTTAATTATAG | |
| pACYC_ $P_{BAD}$_*pxaA*_ R | JE_G_for3.1 | GTTTTTTTGGGCTAACAGGAGGAATTCCATGAAAACTTCACAATTAGTACCTCTTACC | pFF1_NRPS -1 |
| | JE_G_Rxeop | GCAGCAGCCTAGGTTAATTAATTACTTACTTTCAGGTTTATATGACGGTATGC | |
| pACYC_ $P_{BAD}$_*pxaA*_ T2stop | JE_G_for3.1 | GTTTTTTTGGGCTAACAGGAGGAATTCCATGAAAACTTCACAATTAGTACCTCTTACC | *X. stockiae* gDNA |
| | JE_G_TEstop | GCAGCAGCCTAGGTTAATTAATTAGGGGACAAGCGGACTAAATTTG | |
| | JE_G_for2.1 | GTTTTTTTGGGCTAACAGGAGGAATTCCATGAATATTGCTACTGAACTTGCATTTCG | |

| | | | |
|---|---|---|---|
| pACYC_P$_{BAD}$_*xhpA* | JE_G_rev3.1 | GCAGCAGCCTAGGTTAATTAATTACTCATTTTCCAGTACCTCACTTG | *X. hominickii* gDNA |
| pACYC_P$_{BAD}$_*pxaA*_TE-S83A | JE_TE_S83A_for | GTTGGGCATATGGTGGATTTGTCG | *X. stockiae* gDNA |
| | JE_TE_S83A_rev | ACCATATGCCCAACCTCCGATTG | |
| pJE6 | JE_G_for2.1 | GTTTTTTTGGGCTAACAGGAGGAATTCCATGAATATTGCTACTGAACTTGCATTTCG | pACYC_*xhpA* |
| | je0029 | AAGAACATCTTTTTTTGTACAGAGCGCACG | |
| | je0028 | CCGTGCGCTCTGTACAAAAAAAGATGTTCTTTATGATGATGCTCAGACTGAAACTGA | pACYC_*pxaA* |
| | ck1008 | CATGGAATTCCTCCTGTTAGCCCAA | |
| pJE8 | JE_G_for2.1 | GTTTTTTTGGGCTAACAGGAGGAATTCCATGAATATTGCTACTGAACTTGCATTTCG | pACYC_*xhpA* |
| | JE_G_rev3.1 | GCAGCAGCCTAGGTTAATTAATTACTCATTTTCCAGTACCTCACTTG | |
| | ck1011 | TTAATTAACCTAGGCTGCTGCCAC | pACYC_*pxaA* |
| | ck1008 | CATGGAATTCCTCCTGTTAGCCCAA | |
| pJE2 | je0021 | GCTCATTGGTAAAAACGGAGGTTTCGGCTATCCCATCAGTTGAGCTGGGAGAGGGT | pACYC_*xhpA* |
| | JE_G_rev3.1 | GCAGCAGCCTAGGTTAATTAATTACTCATTTTCCAGTACCTCACTTG | |
| | ck1011 | TTAATTAACCTAGGCTGCTGCCAC | pACYC_*pxaA* |
| | je0020 | GATAGCCGAAACCTCCGTTTTTACC | |
| pJE4 | je0024 | GAAGCTCAGCGTGCAGGATAATTTTTACACATTAGGTGGAGATTCCATCCTGATGCTGA | pACYC_*xhpA* |
| | JE_G_rev3.1 | GCAGCAGCCTAGGTTAATTAATTACTCATTTTCCAGTACCTCACTTG | |
| | ck1011 | TTAATTAACCTAGGCTGCTGCCAC | pACYC_*pxaA* |
| | je0025 | TAATGTGTAAAAATTATCCTGCACGCTG | |
| pJE5 | ck1011 | TTAATTAACCTAGGCTGCTGCCAC | pACYC_*xhpA* |
| | je0027 | CTCATACTTTTTCAGGAAACTAATCTGTTTCG | |

| | | | |
|---|---|---|---|
| | je0026 | CACACCGAAACAGATTAGTTTCCTGAAAAAGTATGAGCAAGGGCCGCTTACTGAATTCTCA | pACYC_*pxaA* |
| | JE_G_rev5.1 | GCAGCAGCCTAGGTTAATTAATCAATGGACGGGACATATTTTCTCC | |
| pJE1 | je0019 | AGTGGCACGACTTGCTCTTTTCGAGAATGGTCCAATAAAACCTTACAGCC | pACYC_*xhpA* |
| | JE_G_rev3.1 | GCAGCAGCCTAGGTTAATTAATTACTCATTTTCCAGTACCTCACTTG | |
| | ck1011 | TTAATTAACCTAGGCTGCTGCCAC | pACYC_*pxaA* |
| | je0018 | CTCGAAAAGAGCAAGTCGTGCCAC | |
| pJE3 | je0022 | AGATTGAGAAGCTCAGCGTGCAGGATAATTTTTATGTTCTTGGTGGAGATTCCA | pACYC_*xhpA* |
| | JE_G_rev3.1 | GCAGCAGCCTAGGTTAATTAATTACTCATTTTCCAGTACCTCACTTG | |
| | je0023 | ATTATCCTGCACGCTGAGCTTCTCAATC | pACYC_*pxaA* |
| | ck1011 | TTAATTAACCTAGGCTGCTGCCAC | |
| pJE7 | je0031 | ATCGTCGTCAGGATAGATGTTATGTTGG | pACYC_*xhpA* |
| | ck1011 | TTAATTAACCTAGGCTGCTGCCAC | |
| | je0030 | GGCCAACATAACATCTATCCTGACGACGATTTTTACACATTAGGCGGTGA | pACYC_*pxaA* |
| | JE_G_rev5.1 | GCAGCAGCCTAGGTTAATTAATCAATGGACGGGACATATTTTCTCC | |
| pJE14 | JE_G_for2.1 | GTTTTTTTGGGCTAACAGGAGGAATTCCATGAATATTGCTACTGAACTTGCATTTCG | pACYC_*xhpA* |
| | je0045 | CGCCAGCACAGAGATAAATTCCTGAGAAGATATTTCAGGAATGATATTCCCTTGAGCAGA | |
| | je0046 | GAAATATCTTCTCAGGAATTTATCTCTGTGCTG | pACYC_*pxaA* |
| | ck1008 | CATGGAATTCCTCCTGTTAGCCCAA | |
| pJE13 | JE_G_for3 | TTTGGGCTAACAGGAGGAATTCCATGAAAACTTCACAATTAGTACCTC | *X. stockiae* gDNA |
| | je0043 | TCGATTCAAACGGTTCTTCCGACAGAGATTTATTGCTTTGTGTAAGCGGCTGGC | |
| | je0044 | TCTCTGTCGGAAGAACCGTTTGAATC | *X. hominickii* gDNA |
| | je0037 | CACAATTCCACACATTATACGAGCCGATGATTAATTGTTACTCATTTTCCAGTACCTCACTTGCAGC | |
| pJE39 | je0077 | TATCGTGCGCCAAAAGACCAATAC | |

| | | | |
|---|---|---|---|
| | JE_G_rev3.1 | GCAGCAGCCTAGGTTAATTAATTACTCATTTTCCAGTACCTCACTTG | *X. hominickii* gDNA |
| | ck1011 | TTAATTAACCTAGGCTGCTGCCAC | pACYC_*pxaA* |
| | je0078 | GTATTGGTCTTTTGGCGCACGATATTCTGCGTTATTATGGCATTGCATGAC | |
| pJE40 | je0079 | ATTTTCGATTGTGGCGCAACCTCG | *X. hominickii* gDNA |
| | JE_G_rev3.1 | GCAGCAGCCTAGGTTAATTAATTACTCATTTTCCAGTACCTCACTTG | |
| | ck1011 | TTAATTAACCTAGGCTGCTGCCAC | pACYC_*pxaA* |
| | je0080 | CGAGGTTGCGCCACAATCGAAAATATTCTGGTGAACAGATACAGCAGG | |
| pJE41 | je0081 | TATATTGCGCCACGAGACAATTATG | *X. stockiae* gDNA |
| | JE_G_rev5.1 | GCAGCAGCCTAGGTTAATTAATCAATGGACGGGACATATTTTCTCC | |
| | ck1011 | TTAATTAACCTAGGCTGCTGCCAC | pACYC_*xhpA* |
| | je0082 | CATAATTGTCTCGTGGCGCAATATATTCTCTGTTTGACTCACTTCTGATTTTC | |
| pJE42 | je0083 | CTGTTTGATATCGGTGCTACTTCATTAAC | *X. stockiae* gDNA |
| | JE_G_rev5.1 | GCAGCAGCCTAGGTTAATTAATCAATGGACGGGACATATTTTCTCC | |
| | ck1011 | TTAATTAACCTAGGCTGCTGCCAC | pACYC_*xhpA* |
| | je0084 | GTTAATGAAGTAGCACCGATATCAAACAGATTCTGTTCTGGAGAAATTTGGTGAATC | |
| pJE22 | je0039 | TGAGATAGCTGCAGTCAGC | pJW76 |
| | je0038 | CAATTAATCATCGGCTCGTATAATGTG | |
| | je0056 | CCGTTTTTTTGGGCTAACAGGAGGAATTCCATGAAAAATGATAAGGTGATGACTCTGCCA | pACYC_*pxaA* |
| | je0041 | CACAATTCCACACATTATACGAGCCGATGATTAATTGTCAATGGACGGGACATATTTTCTCCG | |
| pJE43 | JE_G_for3.1 | GTTTTTTTGGGCTAACAGGAGGAATTCCATGAAAACTTCACAATTAGTACCTCTTACC | pACYC_*pxaA* |
| | je0142 | TCGATTTTAATTCCTCCTTCTCGTTTTGTTTTGAGAATTCAGTAAGCG | |
| | je0035 | AACGAGAAGGAGGAATTAAAATCGAAAAAGG | pNA10 |

| | | | |
|---|---|---|---|
| | je1008 | CATGGAATTCCTCCTGTTAGCCCAA | |
| pJE44 | JE_G_for3.1 | GTTTTTTTGGGCTAACAGGAGGAATTCCATGAAAACTTCACAATTAGTACCTCTTACC | pACYC_*pxaA* |
| | je0143 | TTCGATTTTAATTCCTCCTTCTCGTTAAATTCCTGAGAAGATATTTCTTTATTGCTTTG | |
| | je0035 | AACGAGAAGGAGGAATTAAAATCGAAAAAGG | pNA10 |
| | je1008 | CTGGTACGTGTCGGTCGCCACGATCATTTCTTTGATCTGGGAGGCGATTC | |
| pJE45 | je0144 | TAACGAGCTGACTGCAGCTATCTCAGAAACCTTATCAACGCTATTTGATACTCAG | pACYC_*pxaA* |
| | je0141 | TGAATATATGGTGCCGAC | |
| | sz18for | CAATTAATCATCGGCTCGTA | pCOLA_SZ18 |
| | sz18rev | TGAGATAGCTGCAGTCAG | |
| pJE46 | je0145 | TAACGAGCTGACTGCAGCTATCTCAATCTCTGTGCTGGCGATGATGG | pACYC_*pxaA* |
| | je0141 | TGAATATATGGTGCCGAC | |
| | sz18for | CAATTAATCATCGGCTCGTA | pCOLA_SZ18 |
| | sz18rev | TGAGATAGCTGCAGTCAG | |
| pJE58 | je0183 | TTGGGCTAACAGGAGGAATTCCATGATGAAAACTTCACAATTAGTACCTCTTACC | *X. stockiae* gDNA |
| | je0177 | CCTCCTTCTCGTTGGATCCAGACCCATAGATAGCCGAAACCTCCGTTTTTA | |
| | je0176 | GGGTCTGGATCCAACGAGAAG | pNA10 |
| | je0175 | CATGGAATTCCTCCTGTTAGCC | |
| pJE59 | je0178 | TGCAGCTATCTCAGGTTCGGGATCAGATGATGCTCAGACTGAAACTGAATATC | *X. stockiae* gDNA |
| | je0182 | TATACGAGCCGATGATTAATTGTCATCAATGGACGGGACATATTTTCTCC | |
| | jw0061 | TGACAATTAATCATCGGCTCG | pNA155 |
| | je0179 | TGATCCCGAACCTGAGATAGCTG | |
| pJE61 | je0181 | TGCAGCTATCTCAGGTTCGGGATCAATTGCGCCACGAGACAATTATGAAC | *X. stockiae* gDNA |
| | je0182 | TATACGAGCCGATGATTAATTGTCATCAATGGACGGGACATATTTTCTCC | |

| | | | |
|---|---|---|---|
| | jw0061 | TGACAATTAATCATCGGCTCG | pNA155 |
| | je0179 | TGATCCCGAACCTGAGATAGCTG | |
| pJE27 | je0064 | CCGTTTTTTTGGGCTAACAGGAGGAATTCCATGAGAACATCTGAAAGCTCGT | *X. bovienii* gDNA |
| | je0058 | TGTTTTGAGAATTCAGTAAGCGGCCCTTGCCAAGTGTGCAGTAAAGTGTGACGT | |
| | je0044 | TCTCTGTCGGAAGAACCGTTTGAATC | *X. stockiae* gDNA |
| | JE_G_rev5.1 | GCAGCAGCCTAGGTTAATTAATCAATGGACGGGACATATTTTCTCC | |
| pJE28 | je0064 | CCGTTTTTTTGGGCTAACAGGAGGAATTCCATGAGAACATCTGAAAGCTCGT | *X. bovienii* gDNA |
| | je0065 | CCAATTGATATTCAGTTTCAGTCTGAGCATCATCATAACCGTCCCGGTTTCCCCACA | |
| | je0060 | TATGATGATGCTCAGACTGAAACTGAATATC | *X. stockiae* gDNA |
| | JE_G_rev5.1 | GCAGCAGCCTAGGTTAATTAATCAATGGACGGGACATATTTTCTCC | |
| pJE29 | je0066 | CCGTTTTTTTGGGCTAACAGGAGGAATTCCATGCCTATGTCGTGCAATCGT | *X.* KJ12.1 gDNA |
| | je0067 | TGTTTTGAGAATTCAGTAAGCGGCCCTTGAAAATCCACCAATATCTTTTGCTGT | |
| | je0055 | CAAGGGCCGCTTACTGAATTCTC | *X. stockiae* gDNA |
| | JE_G_rev5.1 | GCAGCAGCCTAGGTTAATTAATCAATGGACGGGACATATTTTCTCC | |
| pJE30 | je0066 | CCGTTTTTTTGGGCTAACAGGAGGAATTCCATGCCTATGTCGTGCAATCGT | *X.* KJ12.1 gDNA |
| | je0068 | CCAATTGATATTCAGTTTCAGTCTGAGCATCATCATACTCATGCGTGACTACCGCAGA | |
| | je0060 | TATGATGATGCTCAGACTGAAACTGAATATC | *X. stockiae* gDNA |
| | JE_G_rev5.1 | GCAGCAGCCTAGGTTAATTAATCAATGGACGGGACATATTTTCTCC | |
| pJE109 | TxlA_3 | CACCAGACCGATGCGCTGTATCC | pJE28 |
| | AR970rvBB | CTAGTATTTCCCCTCTTTCTCTAGT | |
| | TxlA_7 | AATACTAGAGAAAGAGGGGAAATACTAGATGAATATGACACGTAACCATACATCCTC | *X. indica* gDNA |
| | TxlA_8 | TACAGCGCATCGGTCTGGTGAAAATCTACCAATAGTTTCTGGCGCTC | |
| pJE110 | TxlA_10 | AACGCCACTCAGCAGAACTTCACGC | pJE30 |

| | | | |
|---|---|---|---|
| | AR970rvBB | CTAGTATTTCCCCTCTTTCTCTAGT | |
| | TxlA_7 | AATACTAGAGAAAGAGGGGAAATACTAGATGAATATGACACGTAACCATACATCCTC | *X. indica* gDNA |
| | TxlA_13 | TGCGTGAAGTTCTGCTGAGTGGCGTTAAAATCTACCAATAGTTTCTGGCGCTC | |
| pJE47 | je0146 | TTTTTGGGCTAACAGGAGGAATTCCATGATTTTTCTTGCTGAAAAACAGATCATTC | *X. bovienii* gDNA |
| | je0147 | TTGAGAATTCAGTAAGCGGCCCTTGAAAATCATAGAGAACCTGCCGGCGT | |
| | je0055 | CAAGGGCCGCTTACTGAATTCTC | pACYC_*pxaA* |
| | ck1008 | CATGGAATTCCTCCTGTTAGCCCAA | |
| pJE48 | je0146 | TTTTTGGGCTAACAGGAGGAATTCCATGATTTTTCTTGCTGAAAAACAGATCATTC | *X. bovienii* gDNA |
| | je0148 | CAATTGATATTCAGTTTCAGTCTGAGCATCATCATATTCCCGTGTGAGGACGG | |
| | je0060 | TATGATGATGCTCAGACTGAAACTGAATATC | pACYC_*pxaA* |
| | JE_G_rev5.1 | GCAGCAGCCTAGGTTAATTAATCAATGGACGGGACATATTTTCTCC | |
| pJE49 | je0149 | TTTTTGGGCTAACAGGAGGAATTCCATGAAAGGTAGTATTGCTAAAAAGGGA | *X. szentirmaii* gDNA |
| | je0150 | TTGAGAATTCAGTAAGCGGCCCTTGCCAAGTGGATGTGTTCCTGGTA | |
| | je0055 | CAAGGGCCGCTTACTGAATTCTC | pACYC_*pxaA* |
| | ck1008 | CATGGAATTCCTCCTGTTAGCCCAA | |
| pJE50 | je0149 | TTTTTGGGCTAACAGGAGGAATTCCATGAAAGGTAGTATTGCTAAAAAGGGA | *X. bovienii* gDNA |
| | je0151 | CAGTTTCAGTCTGAGCATCATCATAATGCTGACGGGCAAATGCGTT | |
| | je0060 | TATGATGATGCTCAGACTGAAACTGAATATC | pACYC_*pxaA* |
| | JE_G_rev5.1 | GCAGCAGCCTAGGTTAATTAATCAATGGACGGGACATATTTTCTCC | |
| pJE51 | je0152 | TTTTTGGGCTAACAGGAGGAATTCCATGAGATCATTTGAGGATTCACTGAATTC | *X. innexi* gDNA |
| | je0153 | TTTGAGAATTCAGTAAGCGGCCCTTGCCAGCGATGCAGCAGAGTATG | |
| | je0055 | CAAGGGCCGCTTACTGAATTCTC | pACYC_*pxaA* |
| | ck1008 | CATGGAATTCCTCCTGTTAGCCCAA | |

| | | | |
|---|---|---|---|
| pJE52 | je0152 | TTTTTGGGCTAACAGGAGGAATTCCATGAGATCATTTGAGGATTCACTGAATTC | *X. innexi* gDNA |
| | je0154 | CAGTTTCAGTCTGAGCATCATCATAACTTTCTTTATTGCTGAAGACCGGTT | |
| | je0060 | TATGATGATGCTCAGACTGAAACTGAATATC | pACYC_*pxa A* |
| | JE_G_rev5.1. | GCAGCAGCCTAGGTTAATTAATCAATGGACGGGACATATTTTCTCC | |
| pJE53 | je0155 | TTTTTGGGCTAACAGGAGGAATTCCATGAATATGACACGTAACCATACATCCT | *X. bovienii* gDNA |
| | je0156 | TTGAGAATTCAGTAAGCGGCCCTTGGAAATCCACCAACAGTTGTTGACGT | |
| | je0055 | CAAGGGCCGCTTACTGAATTCTC | pACYC_*pxa A* |
| | ck1008 | CATGGAATTCCTCCTGTTAGCCCAA | |
| pJE54 | je0155 | TTTTTGGGCTAACAGGAGGAATTCCATGAATATGACACGTAACCATACATCCT | *X. indica* gDNA |
| | je0157 | CAGTTTCAGTCTGAGCATCATCATAGCCACGTGTAACAACCGCTGA | |
| | je0060 | TATGATGATGCTCAGACTGAAACTGAATATC | pACYC_*pxa A* |
| | JE_G_rev5.1 | GCAGCAGCCTAGGTTAATTAATCAATGGACGGGACATATTTTCTCC | |
| pJE55 | frs1 | CCGTTTTTTTGGGCTAACAGGAGGAATTCCATGAAAAACAGTGAATCGCCAATCCA | *C. vaccinii* gDNA |
| | je0158 | CAGTTTCAGTCTGAGCATCATCATAATGCGAGCCGCCAAACT | |
| | je0060 | TATGATGATGCTCAGACTGAAACTGAATATC | pACYC_*pxa A* |
| | JE_G_rev5.1. | GCAGCAGCCTAGGTTAATTAATCAATGGACGGGACATATTTTCTCC | |
| pJE56 | je0161 | CAGTTTCAGTCTGAGCATCATCATATTTGGCATGATGATTAATGAAACCTGG | *X. nematophila* gDNA |
| | je0162 | TTTTTGGGCTAACAGGAGGAATTCCATGTTTCTAGATAAAGTCGGGCAGC | |
| | je0055 | CAAGGGCCGCTTACTGAATTCTC | pACYC_*pxa A* |
| | ck1008 | CATGGAATTCCTCCTGTTAGCCCAA | |
| pJE57 | je0161 | CAGTTTCAGTCTGAGCATCATCATATTTGGCATGATGATTAATGAAACCTGG | *X. nematophila* |
| | je0162 | TTTTTGGGCTAACAGGAGGAATTCCATGTTTCTAGATAAAGTCGGGCAGC | |
| | je0055 | CAAGGGCCGCTTACTGAATTCTC | |

| | | | |
|---|---|---|---|
| | ck1008 | CATGGAATTCCTCCTGTTAGCCCAA | pACYC_*pxa A* |
| pJE_frs5 | frs13 | CCGTTTTTTTGGGCTAACAGGAGGAATTCCATGACCGTATCCGATAACGTATTCCTGCG | *C. vaccinii* gDNA |
| | frs14 | GCGGTGGCAGCAGCCTAGGTTAATTAACTACAGCAGCATGGTTTGGCAGC | |
| | ck1008 | CATGGAATTCCTCCTGTTAGCCCAA | pCK_0432 |
| | ck1011 | TTAATTAACCTAGGCTGCTGCCAC | |
| pJE_frs11 | frs17 | CCGTTTTTTTGGGCTAACAGGAGGAATTCCATGAGCAATCCCTTTGATGATAAAGATGGT | *C. vaccinii* gDNA |
| | frs29 | GCGGTGGCAGCAGCCTAGGTTAATTAATCATTTATCATCGCACTCCATTGCA | |
| | ck1008 | CATGGAATTCCTCCTGTTAGCCCAA | pCK_0433 |
| | ck1011 | TTAATTAACCTAGGCTGCTGCCAC | |
| pJE23 | Je0055 | CAAGGGCCGCTTACTGAATTCTC | pACYC_*pxa A* |
| | ck1008 | CATGGAATTCCTCCTGTTAGCCCAA | |
| | je0056 | CCGTTTTTTTGGGCTAACAGGAGGAATTCCATGAAAAATGATAAGGTGATGACTCTGCCA | *X. miraniensis* gDNA |
| | je0057 | TGTTTTGAGAATTCAGTAAGCGGCCCTTGCCATTGATTTAGCAATAGCGCTCGT | |
| pJE24 | je0060 | TATGATGATGCTCAGACTGAAACTGAATATC | pACYC_*pxa A* |
| | ck1008 | CATGGAATTCCTCCTGTTAGCCCAA | |
| | je0056 | CCGTTTTTTTGGGCTAACAGGAGGAATTCCATGAAAAATGATAAGGTGATGACTCTGCCA | *X. miraniensis* gDNA |
| | je0059 | CCAATTGATATTCAGTTTCAGTCTGAGCATCATCATAATACTGGCGTTGATAAGCGGTGC | |
| pJE25 | je0055 | CAAGGGCCGCTTACTGAATTCTC | pACYC_*pxa A* |
| | ck1008 | CATGGAATTCCTCCTGTTAGCCCAA | |
| | je0062 | TGTTTTGAGAATTCAGTAAGCGGCCCTTGGAAATTACAGAGAATTTGTTGACGT | *X. indica* gDNA |
| | je0061 | CCGTTTTTTTGGGCTAACAGGAGGAATTCCATGAAACTATTTTCCTTATCTGAAGC | |
| pJE26 | je0060 | TATGATGATGCTCAGACTGAAACTGAATATC | |

| | | | |
|---|---|---|---|
| | ck1008 | CATGGAATTCCTCCTGTTAGCCCAA | pACYC_*pxaA* |
| | je0063 | CCAATTGATATTCAGTTTCAGTCTGAGCATCATCATATTCTTGTGTGATTACTGCTGAATGATCGGG | *X. indica* gDNA |
| | je0061 | CCGTTTTTTTGGGCTAACAGGAGGAATTCCATGAAACTATTTTCCTTATCTGAAGC | |
| pJE76 | TxlA_4 | AGAGAAAGAGGGGAAATACTAGATGAATAAGGATGGATATTATAATTTAAC | *P. temperate* gDNA |
| | TxlA_5 | GATACAGCGCATCGGTCTGGTGTTTATTATTAATAATCTCTAAATTTCTATATTC | |
| | TxlA_3 | CACCAGACCGATGCGCTGTATCC | pJE28 |
| | AR970rvBB | CTAGTATTTCCCCTCTTTCTCTAGT | |
| pJE80 | TxlA_4 | AGAGAAAGAGGGGAAATACTAGATGAATAAGGATGGATATTATAATTTAAC | *P. temperate* gDNA |
| | TxlA_11 | GCGTGAAGTTCTGCTGAGTGGCGTTTTTATTATTAATAATCTCTAAATTTCTATATTC | |
| | TxlA_10 | AACGCCACTCAGCAGAACTTCACGC | pJE30 |
| | AR970rvBB | CTAGTATTTCCCCTCTTTCTCTAGT | |
| pJE75 | TxlA_1 | ACTAGAGAAAGAGGGGAAATACTAGATGATTTTTCTTGCTGAAAAACAGATC | *X. bovienii* gDNA |
| | TxlA_2 | ATACAGCGCATCGGTCTGGTGAAAATCATAGAGAACCTGC | |
| | TxlA_3 | CACCAGACCGATGCGCTGTATCC | pJE28 |
| | AR970rvBB | CTAGTATTTCCCCTCTTTCTCTAGT | |
| pJE79 | TxlA_1 | ACTAGAGAAAGAGGGGAAATACTAGATGATTTTTCTTGCTGAAAAACAGATC | *X. bovienii* gDNA |
| | TxlA_9 | TGCGTGAAGTTCTGCTGAGTGGCGTTAAAATCATAGAGAACCTGC | |
| | TxlA_10 | AACGCCACTCAGCAGAACTTCACGC | pJE30 |
| | AR970rvBB | CTAGTATTTCCCCTCTTTCTCTAGT | |
| pJE77 | pxaA_SEVA | CTAGAGAAAGAGGGGAAATACTAGATGAAAACTTCACAATTAGTACCTCTTAC | pACYC_*pxaA* |
| | TxlA_6 | CGGATACAGCGCATCGGTCTGGTGGAAAAGAGCAAGTCGTGCCAC | |
| | TxlA_3 | CACCAGACCGATGCGCTGTATCC | pJE28 |

| | | | |
|---|---|---|---|
| | AR970rvBB | CTAGTATTTCCCCTCTTTCTCTAGT | |
| pJE81 | TxlA_12 | CGTGAAGTTCTGCTGAGTGGCGTTGAAAAGAGCAAGTCGTGCCAC | pACYC_*pxaA* |
| | pxaA_SEVA | CTAGAGAAAGAGGGGAAATACTAGATGAAAACTTCACAATTAGTACCTCTTAC | |
| | TxlA_10 | AACGCCACTCAGCAGAACTTCACGC | pJE30 |
| | AR970rvBB | CTAGTATTTCCCCTCTTTCTCTAGT | |
| pJE86 | Je0064 | CCGTTTTTTTGGGCTAACAGGAGGAATTCCATGAGAACATCTGAAAGCTCGT | *X. bovienii* gDNA |
| | Je0228 | GTGAAGTTCTGCTGAGTGGCGTTCCAGCGGTGCAGCAGG | |
| | Ck1008 | CATGGAATTCCTCCTGTTAGCCCAA | pJE30 |
| | TxlA_10 | AACGCCACTCAGCAGAACTTCACGC | |
| pJE87 | Je0152 | TTTTTGGGCTAACAGGAGGAATTCCATGAGATCATTTGAGGATTCACTGAATTC | *X. innexi* gDNA |
| | Je0229 | GGATACAGCGCATCGGTCTGGTGCCAGCGATGCAGCAGAGTATG | |
| | TxlA_3 | CACCAGACCGATGCGCTGTATCC | pJE28 |
| | AR970rvBB | CTAGTATTTCCCCTCTTTCTCTAGT | |
| pJE88 | Je0152 | TTTTTGGGCTAACAGGAGGAATTCCATGAGATCATTTGAGGATTCACTGAATTC | *X. innexi* gDNA |
| | Je0230 | AGTTCTGCTGAGTGGCGTTCCAGCGATGCAGCAGAGTATG | |
| | TxlA_10 | AACGCCACTCAGCAGAACTTCACGC | pJE30 |
| | AR970rvBB | CTAGTATTTCCCCTCTTTCTCTAGT | |
| pJE89 | Je0231 | TTTTGGGCTAACAGGAGGAATTCCATGAATACACAGCGTAACCACAATCC | *X. stockiae* gDNA |
| | Je0232 | GCGGATACAGCGCATCGGTCTGGTGGAAATTTTCCAACAATAGCTGGTGTTC | |
| | TxlA_3 | CACCAGACCGATGCGCTGTATCC | pJE28 |
| | AR970rvBB | CTAGTATTTCCCCTCTTTCTCTAGT | |
| pJE90 | Je0231 | TTTTGGGCTAACAGGAGGAATTCCATGAATACACAGCGTAACCACAATCC | *X. stockiae* gDNA |
| | Je0233 | GCGTGAAGTTCTGCTGAGTGGCGTTGAAATTTTCCAACAATAGCTGGTGTTC | |

| | | | |
|---|---|---|---|
| | TxlA_10 | AACGCCACTCAGCAGAACTTCACGC | pJE30 |
| | AR970rvBB | CTAGTATTTCCCCTCTTTCTCTAGT | |
| pJE91 | Je0064 | CCGTTTTTTTGGGCTAACAGGAGGAATTCCATGAGAACATCTGAAAGCTCGT | pCK_0760 |
| | Je0229 | GGATACAGCGCATCGGTCTGGTGCCAGCGATGCAGCAGAGTATG | |
| | TxlA_3 | CACCAGACCGATGCGCTGTATCC | pJE28 |
| | AR970rvBB | CTAGTATTTCCCCTCTTTCTCTAGT | |
| pJE92 | Je0064 | CCGTTTTTTTGGGCTAACAGGAGGAATTCCATGAGAACATCTGAAAGCTCGT | pCK_0760 |
| | Je0230 | AGTTCTGCTGAGTGGCGTTCCAGCGATGCAGCAGAGTATG | |
| | TxlA_10 | AACGCCACTCAGCAGAACTTCACGC | pJE30 |
| | AR970rvBB | CTAGTATTTCCCCTCTTTCTCTAGT | |
| pJE64 | je0146 | TTTTTGGGCTAACAGGAGGAATTCCATGATTTTTCTTGCTGAAAAACAGATCATTC | *X. indica* gDNA |
| | je0190 | CCTCCTTCTCGTTGGATCCAGACCCAAAATCATAGAGAACCTGCCGGCGT | |
| | je0176 | GGGTCTGGATCCAACGAGAAG | pNA10 |
| | ck1008 | CATGGAATTCCTCCTGTTAGCCCAA | |
| pJE65 | je0155 | TTTTTGGGCTAACAGGAGGAATTCCATGAATATGACACGTAACCATACATCCT | *X. indica* gDNA |
| | je0193 | CCTCCTTCTCGTTGGATCCAGACCCAAAATCTACCAATAGTTTCTGGCGCT | |
| | je0176 | GGGTCTGGATCCAACGAGAAG | pNA10 |
| | ck1008 | CATGGAATTCCTCCTGTTAGCCCAA | |
| pJE66 | je0152 | TTTTTGGGCTAACAGGAGGAATTCCATGAGATCATTTGAGGATTCACTGAATTC | *X. innexi* gDNA |
| | je0192 | CCTCCTTCTCGTTGGATCCAGACCCCCAGATATTCAATAAGGTGTGACG | |
| | je0176 | GGGTCTGGATCCAACGAGAAG | pNA10 |
| | ck1008 | CATGGAATTCCTCCTGTTAGCCCAA | |
| pJE67 | je0149 | TTTTTGGGCTAACAGGAGGAATTCCATGAAAGGTAGTATTGCTAAAAAGGGA | |

| | | | |
|---|---|---|---|
| | je0191 | CCTCCTTCTCGTTGGATCCAGACCCCCAAGTTTCCAGCAGCAATGTAC | *X. szentirmaii* gDNA |
| | je0176 | GGGTCTGGATCCAACGAGAAG | pNA10 |
| | ck1008 | CATGGAATTCCTCCTGTTAGCCCAA | |
| pJE68 | je0112 | CCCGTTTTTTTGGGCTAACAGGAGGAATTCCATGAATAAGGATGGATATTATAATTTAACATC | *P. temperate* gDNA |
| | je0194 | CCTCCTTCTCGTTGGATCCAGACCCTTTATTATTAATAATCTCTAAATTTCTATATTC | |
| | je0176 | GGGTCTGGATCCAACGAGAAG | pNA10 |
| | ck1008 | CATGGAATTCCTCCTGTTAGCCCAA | |

### 3. Supplementary Figures

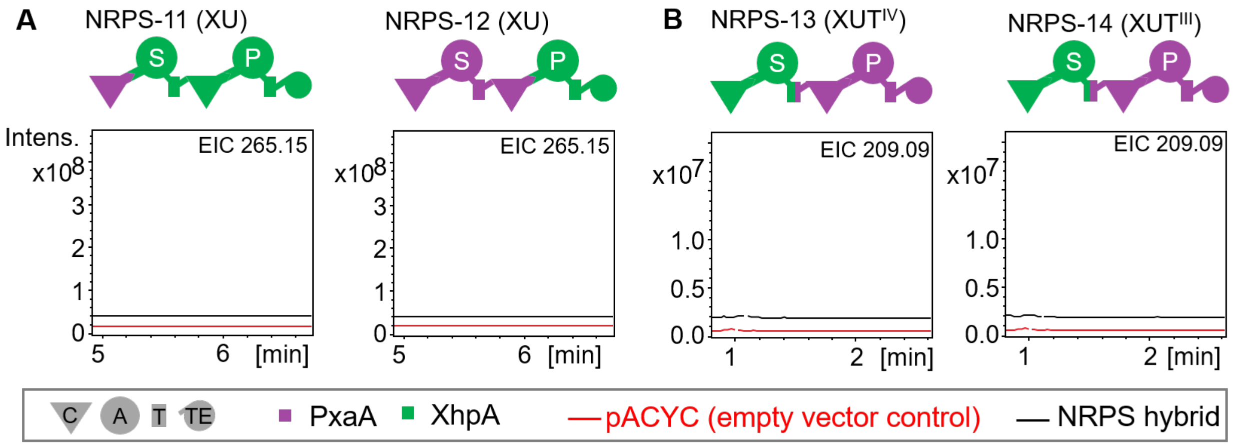


**Figure S1.** NRPS engineering of PxaA by exploiting XhpA.[1] **A**) Engineering PxaA−XhpA hybrids by testing the indicated XU concepts. The EICs of **2** (*m/z* = 265.15 $[M+H]^+$) for the respective NRPS hybrids (black) were compared to the empty vector control (red). **B**) Engineering XhpA−PxaA hybrids by testing the indicated XU approaches. The EICs of **5** (*m/z* = 209.09 $[M+H]^+$) for the respective NRPS hybrids (black) were compared to the empty vector control (red). Peak intensity of the parental PxaA and XhpA variant were 4 x $10^8$ and 1 x $10^7$, respectively.

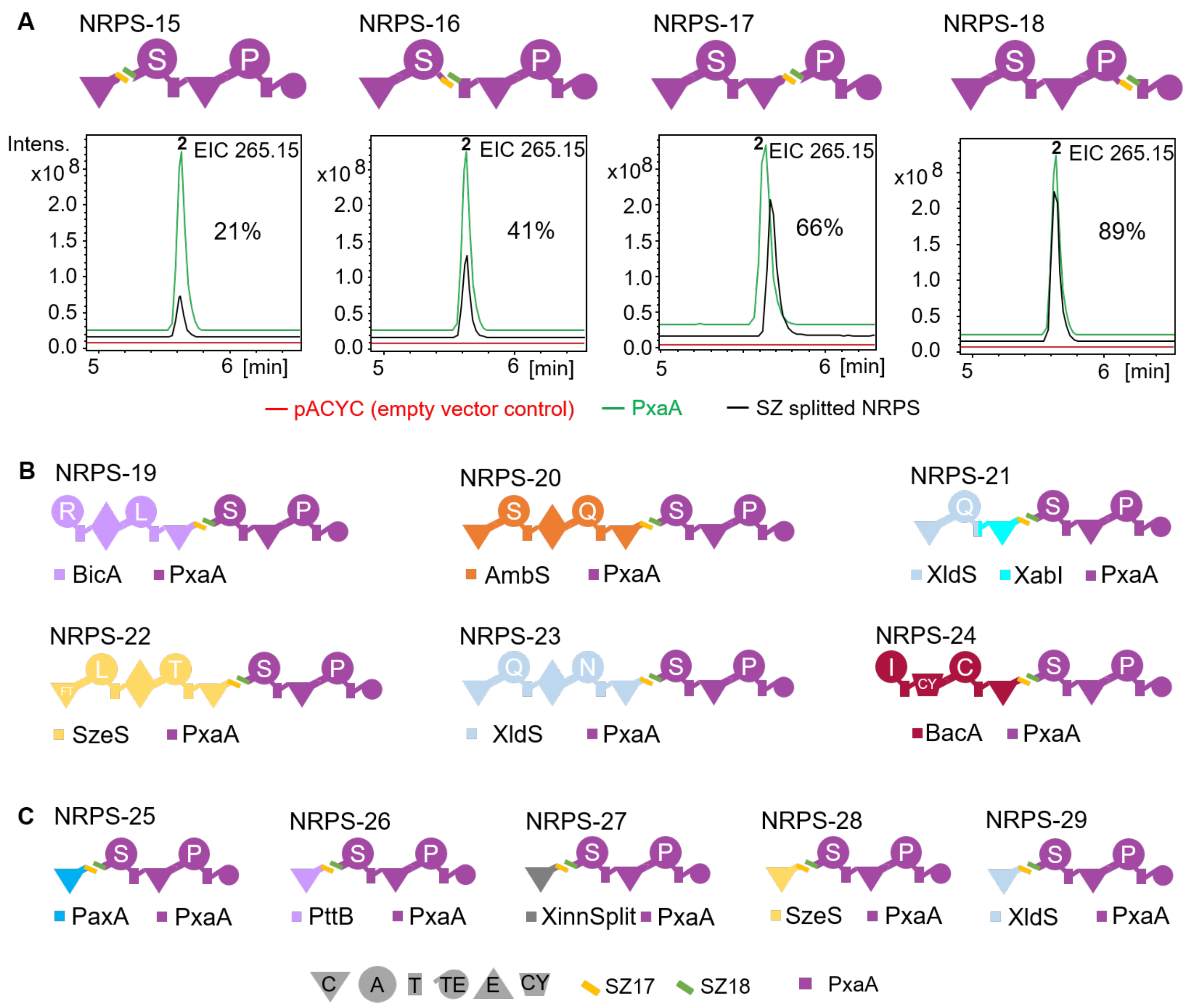


**Figure S2.** Generating SZ17/18-split PxaA variants. **A**) SYNZIPs SZ17:18[2,9] were introduced between $C_1$-$A_1$, $A_1$-$T_1$, $C_2$-$A_2$, and $A_2$-$T_2$ linker. The EICs of **2** (*m/z* = 265.15 $[M+H]^+$) for the NRPS hybrids (black) were compared to the empty vector control (red), and to the wild-type PxaA (green; Fig. 2). Relative quantification was performed for the production of **2** by the bi-partite PxaA (given in %). For relative quantification, the production of the WT PxaA was set to 100%. **B**) SZ18 was introduced into the PxaA $C_1$-$A_1$ linker. A set of available SZ17-NRPSs[8] were used and co-expressed with SZ18-PxaA. **C**) A set of plasmids encoding $C_{starter}$ domains including SZ17 was created and co-expressed with SZ18-PxaA.

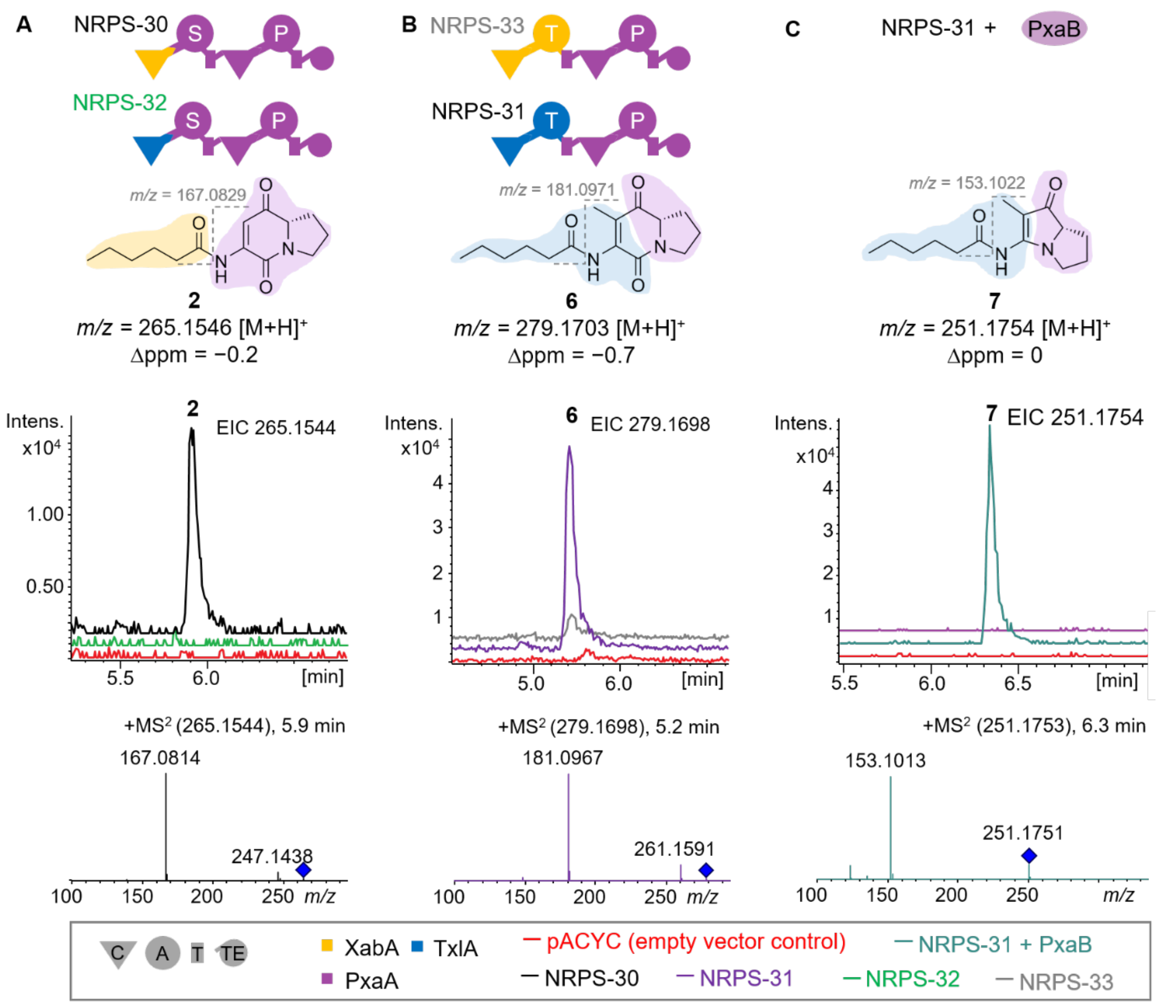


**Figure S3.** Engineering of functional NRPS hybrids XabA-PxaA and TxlA-PxaA. **A**) The $C_{starter}$ domain of the TxlA NRPS and the XabA NRPS were introduced using the XU concept into PxaA, yielding NRPS-30 and NRPS-32, respectively. NRPS-30 produced **2** (black chromatogram), as judged by comparison to the empty vector control (red chromatogram). The proposed structure of **2** is based on LC-HRMS/MS analysis. EIC for **2** (*m/z* calc'd for $C_{14}H_{20}N_2O_3$ $[M+H]^+$ = 265.1547; obs'd. *m/z* = 265.1544; Δppm = −0.2). Inset: proposed structure of **2** indicating the position of $MS^2$ fragmentation (*m/z* calc'd for $C_8H_{10}N_2O_2$ $[M+H]^+$ = 167.0815). **B**) The $C_{starter}$–A domains of the TxlA NRPS and the XabA NRPS were introduced into PxaA using the XUT concept, yielding NRPS-33 and NRPS-31, respectively. Both NRPS-33 and NRPS-31 produced **6**, as judged by comparison to the empty vector control. The proposed structure of **6** is based on LC-HRMS/MS analysis. Inset: proposed structure of **6** (*m/z* calc'd for $C_{15}H_{22}N_2O_3$ $[M+H]^+$ = 279.1703; obs'd. *m/z* = 279.1698; Δppm = −0.7). The proposed structure for the key observed $MS^2$ fragment is indicated (*m/z* calc'd for $C_9H_{12}N_2O_2$ $[M+H]^+$ = 181.0971). **C**) The identity of **7** was proposed based on LC-HRMS/MS analysis. Inset: proposed structure of **7** (*m/z* calc'd for $C_{14}H_{22}N_2O_2$ $[M+H]^+$ = 251.1754; obs'd. *m/z* = 251.1754; Δppm = 0). The key $MS^2$ fragment is indicated (*m/z* calc'd for $C_8H_{12}N_2O$ $[M+H]^+$ = 153.1022). Blue diamonds indicate the parent ion.

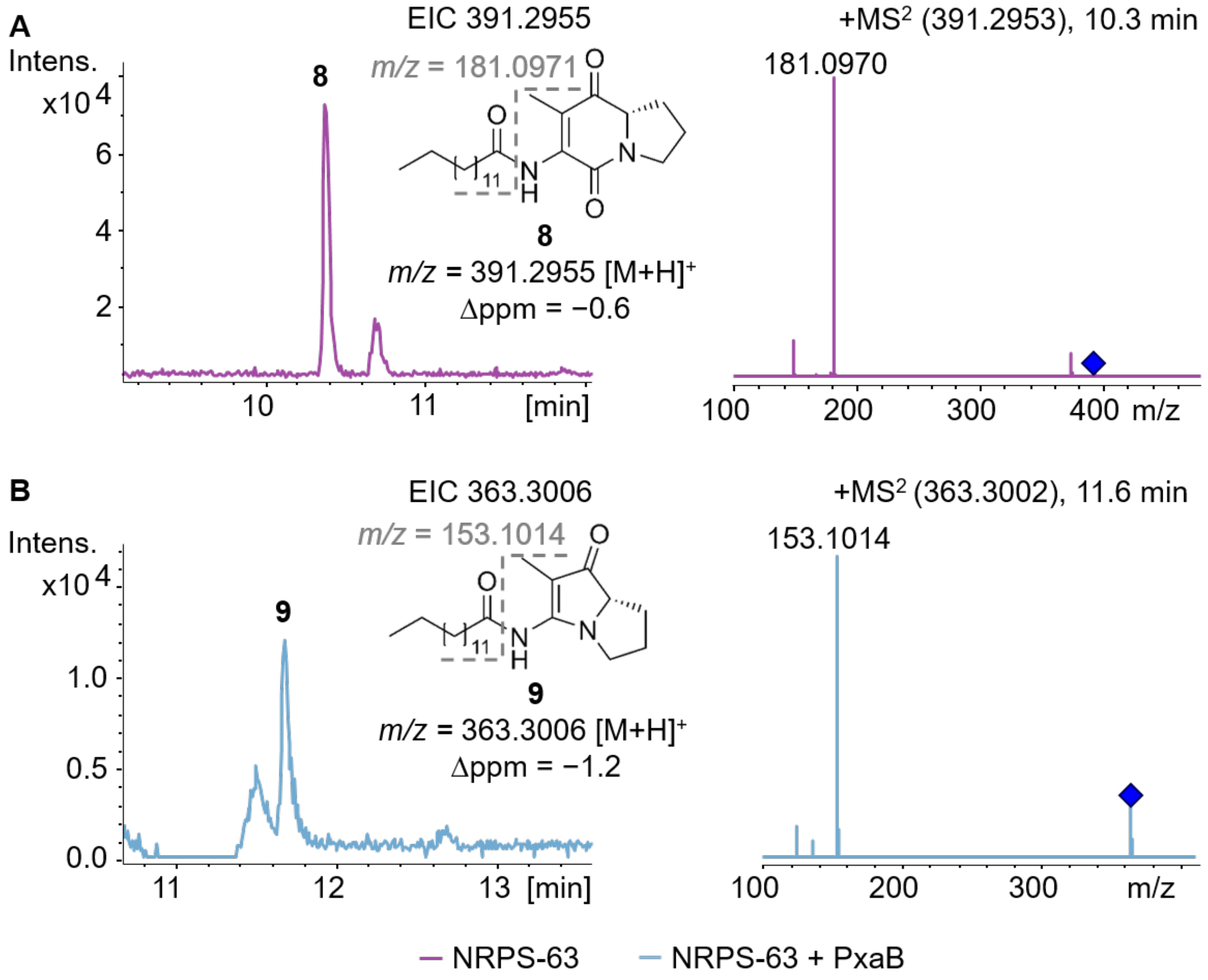


**Figure S4.** The $C_{starter}$ domain of the XldS NRPS was introduced into NRPS-33 and -31 (See Fig. S3) using the XU concept, yielding NRPS-63 and NRPS-64, respectively. **A**) The proposed structure of **8** is based on LC-HRMS/MS analysis. Inset: proposed structure of **8** (*m/z* calc'd for $C_{23}H_{38}N_2O_3$ $[M+H]^+$ = 391.2955; obs'd. *m/z* = 391.2953; Δppm = −0.6). Proposed structure for the key observed $MS^2$ fragment is indicated (*m/z* calc'd for $C_9H_{12}N_2O_2$ $[M+H]^+$ = 181.0971). **B**) Co-expression of both NRPS-63 and NRPS-64 with PxaB led to the production of **9**. The structure of **9** was proposed by LC-HRMS/MS analysis. Inset: proposed structure of **9** (*m/z* calc'd for $C_{22}H_{38}N_2O_2$ $[M+H]^+$ = 363.3006; obs'd. *m/z* = 363.3002; Δppm = −1.2). Proposed structure for the key observed $MS^2$ fragment is indicated (*m/z* calc'd for $C_8H_{12}N_2O$ $[M+H]^+$ = 153.1022). Blue diamonds indicate the parent ion.

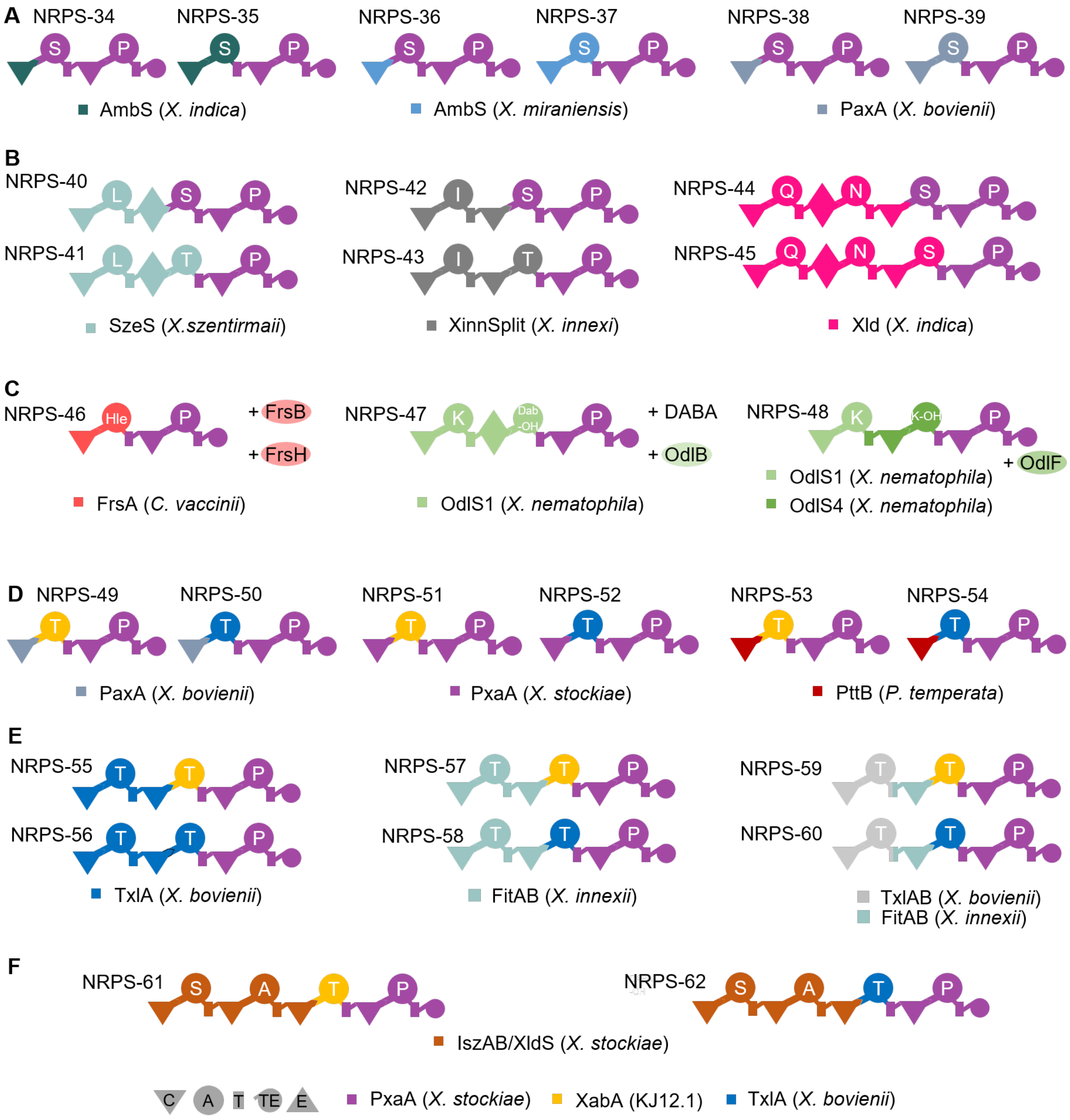


**Figure S5.** Further engineering of PxaA (**A**–**C**) and NRPS-30 and NRPS-31 (**D**–**F**) led to a set of non-functional NRPS hybrids. In the case of hybrids shown in **C**), additional enzymes were co-expressed that are required for the production of the NRPS substrates. FrsB = MbtH-like protein,[10] FrsH = non-heme monooxygenase,[10] OdlB/OdlF = hydroxylases[6]. The second module in OdlS1 is known to incorporate Dab-OH (3-hydroxy-2,4-diaminobutyric acid), while OdlS4 is known to incorporate K-OH (δ-hydroxy lysine).

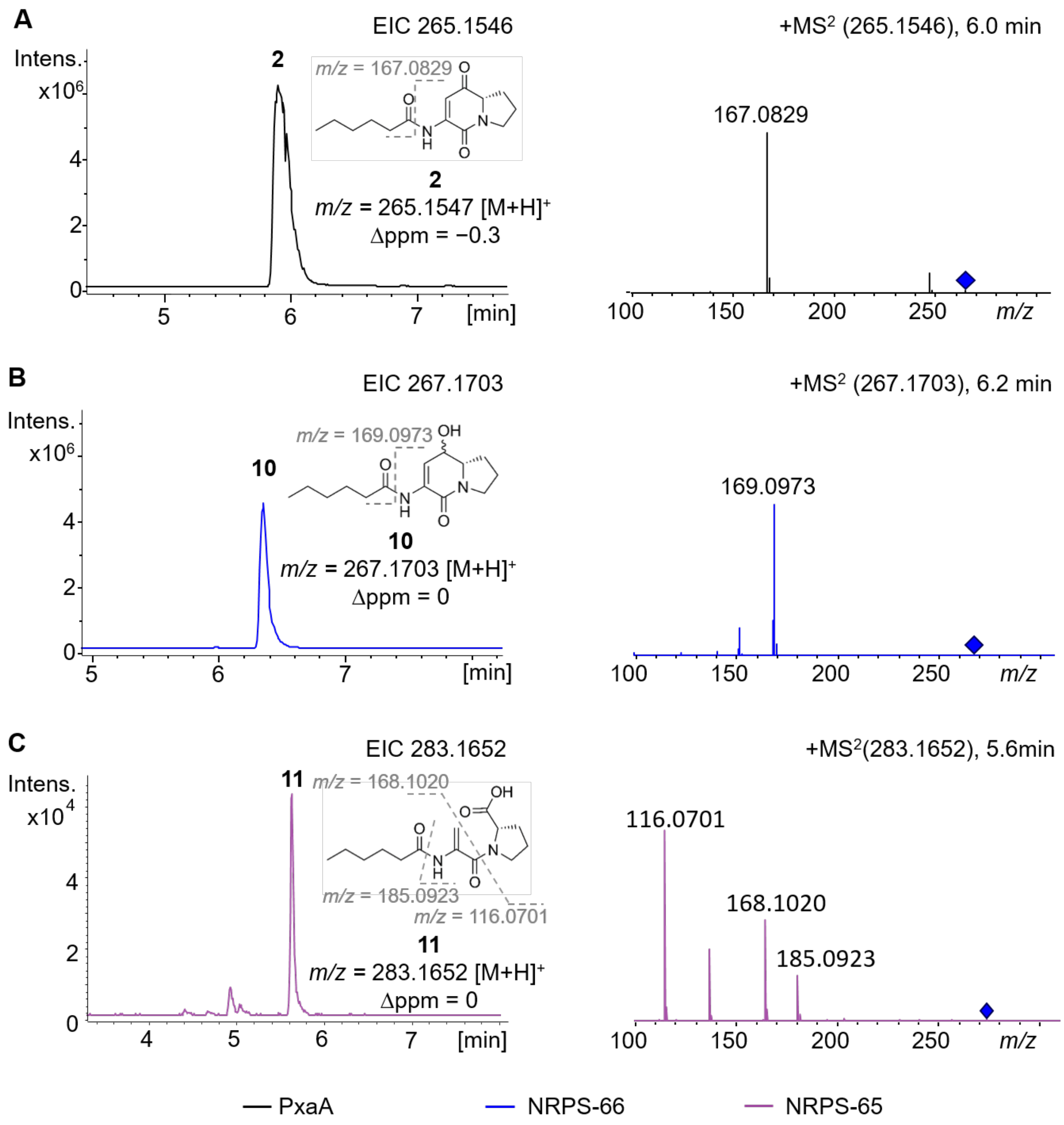


**Figure S6.** Investigation of the role of the TE domain in PxaA. **A**) PxaA produces **2** as judged by LC-HRMS/MS analysis. The structure of **2** (*m*/*z* calc'd for $C_{14}H_{20}N_2O_3$ $[M+H]^+$ = 265.1547; obs'd *m/z* = 265.1546; Δppm = −0.3) is shown. Inset: proposed structure for the key observed $MS^2$ fragment (*m*/*z* calc'd for $C_8H_{10}N_2O_2$ $[M+H]^+$ = 167.0815). **B**) NRPS-66 produces **11** as judged by LC-HRMS/MS analysis. The structure of **11** (*m*/*z* calc'd for $C_{14}H_{22}N_2O_3$ $[M+H]^+$ = 267.1703; obs'd *m/z* = 267.1703; Δppm = 0) is shown. Inset: proposed structure for the key observed $MS^2$ fragment (*m*/*z* calc'd for $C_8H_{12}N_2O_2$ $[M+H]^+$ = 169.0971). **C**) NRPS-65 produces **10** as judged by LC-HRMS/MS analysis. The structure of **10** (*m*/*z* calc'd for $C_{14}H_{22}N_2O_4$ $[M+H]^+$ = 283.1652; obs'd *m/z* = 283.1652; Δppm = 0) is shown. Inset: proposed structure for the key observed $MS^2$ fragment (*m*/*z* calc'd for $C_8H_{12}N_2O_3$ $[M+H]^+$ = 185.0920; *m*/*z* calc'd for $C_5H_9NO_2$ $[M+H]^+$ = 116.0706; *m*/*z* calc'd for $C_9H_{14}NO_2$ $[M+H]^+$ =168.2129). Blue diamonds indicate the parent ion.

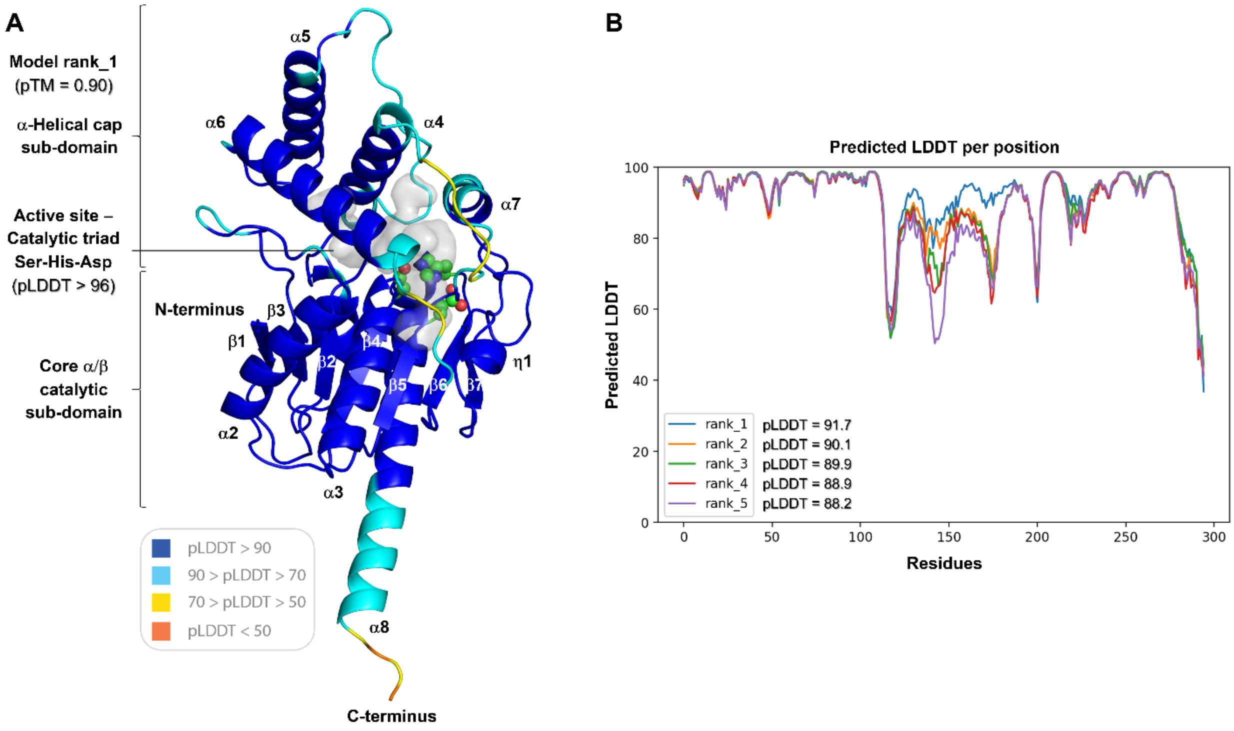


**Figure S7.** AlphaFold2[3]-derived structural model of the PxaA TE domain. **A**) The thioesterase domain belongs to the α/β hydrolase family. Members of this family catalyze a broad range of reactions and share a characteristic fold comprising a catalytic core and a prominent α-helical subdomain.[1] Concerning the selected model (rank_1), it exhibits a high-confidence score (predicted Template Modeling (pTM) = 0.90)), attesting to the prediction accuracy of both the overall fold and the relative arrangement of the subdomains**.** The active site Ser-His-Asp catalytic triad (shown in sphere form) is situated at the interdomain interface, and exhibits a predicted Local Distance Difference Test (pLDDT) > 96, meaning that the side chain positions have been predicted with high confidence. **B**) Comparison among the five AlphaFold2-derived structures shows that the rank_1 model is clearly superior to the others (pLDDT = 91.7), with differences apparent in the α-helical cap subdomain.

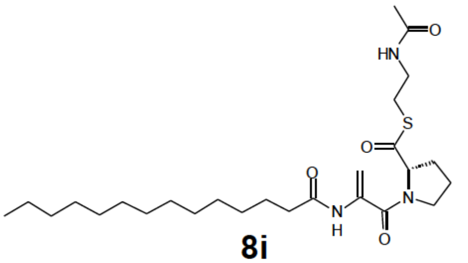

**8i**

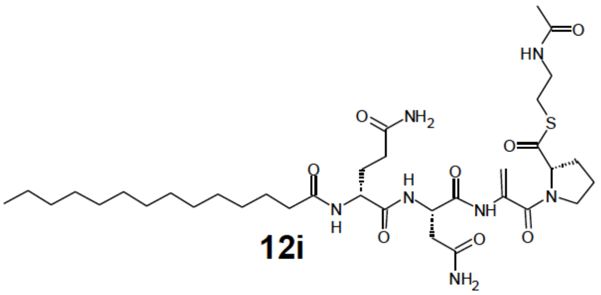

**12i**

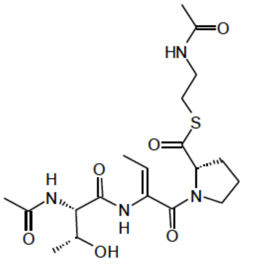

**13i**

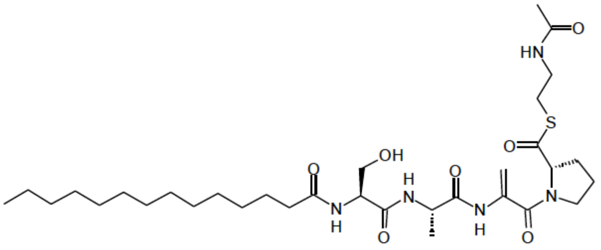

**14i**

**Figure S8.** Structures of SNAC-peptides **8i**, **12i**-1**4i**, predicted model intermediates of NRPS-63/-64 (**8i**), NRPS-44/-45 (**12i**), NRPS-57/58 (**13i**) and NRPS-61/-62 (**14i**) (for the respective NRPS structures see Fig. S5).
